# On the limits of detection of epistatic higher-order interactions

**DOI:** 10.64898/2026.03.05.709494

**Authors:** José Camacho-Mateu, Giulio Burgio, Isabel Quirós-Rodriguez, Miguel D. Fernandez-de-Bobadilla, Alvaro Sanchez

**Affiliations:** Institute of Functional Biology and Genomics IBFG-CSIC, University of Salamanca, 37007 Salamanca, Spain

## Abstract

The function of microbial communities is often dominated by additive and pairwise interactions, raising the question of whether this reflects intrinsic biological simplicity or fundamental limits of detection. Here, we leverage the theory of fitness landscapes to bridge microbial ecology and genetics, and show that this apparent simplicity is a generic consequence of structural and statistical constraints rather than evidence for intrinsically weak higher-order interactions (HOIs). We separate the detectability of individual epistatic interactions from their contribution to functional variance, and demonstrate that local *k*-order interactions suffer from exponential noise amplification while their contributions to total variance are intrinsically suppressed by combinatorial geometric dilution. Applying this framework to a fully sampled 2^10^ experimental microbial landscape, we find that only first- and second-order interactions are distinguishable from experimental noise. Furthermore, generalized Lotka-Volterra simulations reveal that experimental noise alone can generate the illusion of higher-order structure in communities where all direct mechanistic interactions are pairwise and indirect interactions are weak. Our findings identify universal, order-dependent limits on the quantification of epistasis that apply to high-dimensional landscapes across ecology and genetics, providing a principled foundation for rational community design.

## I. INTRODUCTION

Microbial communities are essential drivers of global biogeochemical cycles [1] and crucial components of human health [2] and industrial biotechnology [3]. Predicting and engineering the function of these complex systems–such as total biomass or metabolite production– requires a rigorous understanding of how community composition translates into collective performance [4–7].

Inspired by the concept of fitness landscapes in genetics, an *ecological landscape*, or *community-function landscape* [8–12], describes how a community-level function depends on the presence or absence of individual species. In this framework, species play the role of genetic loci, and each possible community configuration corresponds to a point in a high-dimensional binary space. The landscape assigns a quantitative functional value to every combination of species, thereby defining a discrete map from community composition to ecosystem-level outcome (see Fig. 1A and B).

**FIG. 1.**
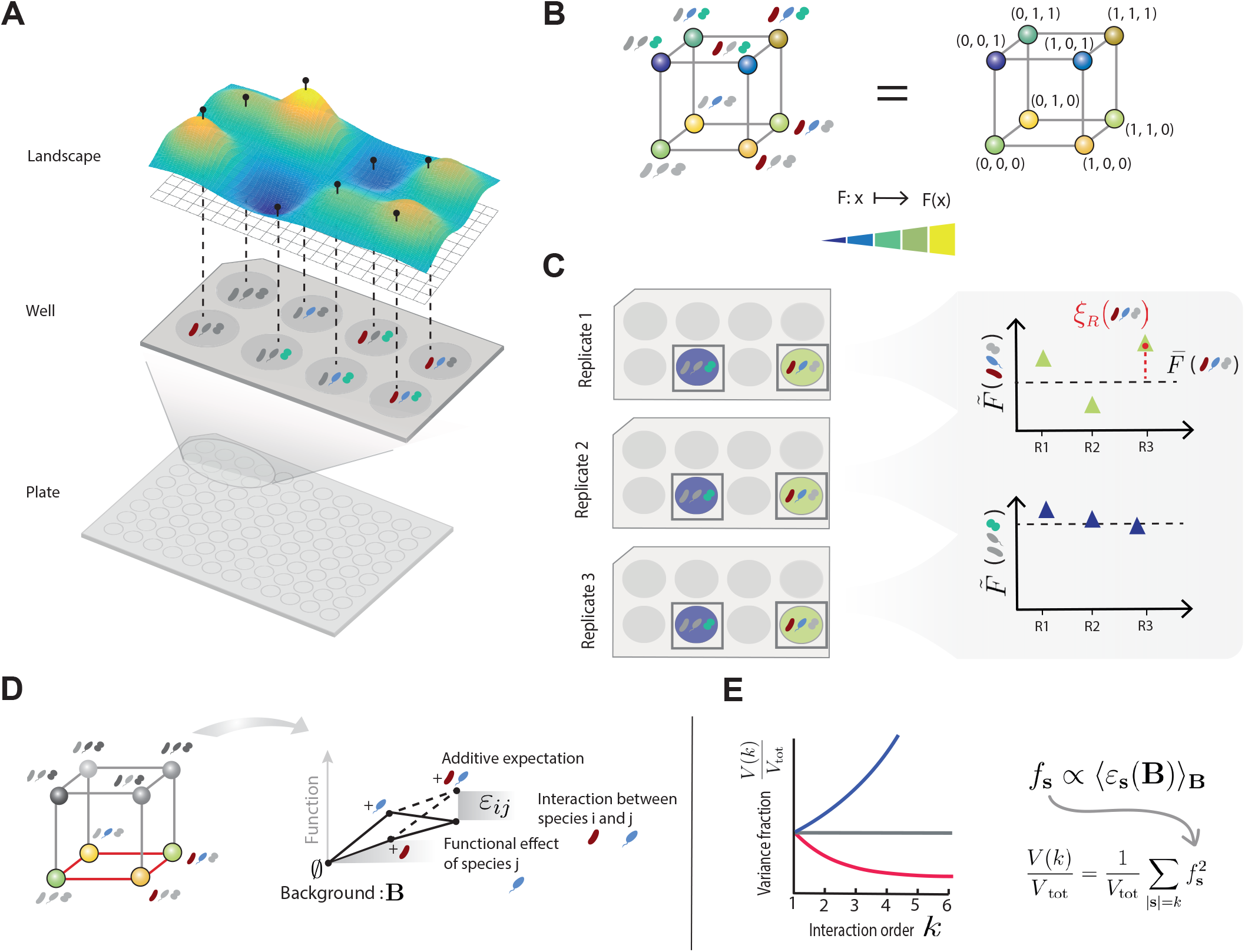
Quantifying the detectability of high-order interactions in community-function landscapes. **(A)** An ecological (community–function) landscape maps the set of all possible community compositions, defined by species presence or absence, to a collective community-level function *F*. The panel illustrates this concept for a three-species example and shows how community compositions are experimentally sampled using a 96-well plate. **(B)** Beyond the metaphorical depiction in (A), the community-function landscape is rigorously defined by a function *F* on the binary hypercube {0, 1} ^*N*^, where each vertex **x** represents a presence/absence community composition. The color of each vertex encodes the value of *F*(**x**). **(C)** The underlying biological signal 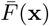, representing the true community-level function, is not directly observable and is instead measured through a finite number of noisy experimental observations 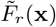. After correcting for batch-to-batch effects, the remaining variability corresponds to stochastic residual fluctuations *ξ*_*r*_ (**x**) around a true biological signal. **(D)** Ecological epistasis *ε*_*ij*_ (**B**) defines a functional interaction and is measured as the deviation from additivity (dashed lines) when focal species are added to a fixed background **B. (E)** The order-resolved variance spectrum 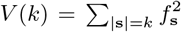 quantifies the contribution of each interaction order to the total variance in the community-function landscape. The panel illustrates three possible scenarios: (blue) variance that increases with interaction order, (gray) all orders contribute similarly to the variance, and (red) the variance decreases with order. The central question we ask in this paper is whether the often reported dominance of low-order interactions in the variance spectrum reflects genuine biological simplicity or the masking of higher-order contributions by either the combinatorial amplification of noise or the rarefied combinatorial sampling of high-order interactions in landscapes of small size (*N <* 10 species).

This composition-to-function mapping is inherently non-linear and structured by interactions among species [6]. While simple pairwise interactions are often the focus of ecological models [13, 14], recent work suggests that higher-order interactions (HOIs)–involving three or more species simultaneously–can play an important role in shaping community function [15–18]. In particular, mechanistic studies of microbial communities have shown that HOIs can emerge naturally from basic ecological processes such as resource competition and metabolic crossfeeding [6]. However, the problem with higher-order interactions is not their existence, but their detectability under noisy and limited replication conditions. The combinatorial structure of ecological landscapes implies that resolving an interaction of order *k* requires probing on the order of 2^*k*^ distinct community configurations in presence–absence space, each associated with a functional measurement. As a result, higher-order contributions may be strongly confounded by statistical uncertainty, raising the question of which interaction orders can be reliably inferred from data and under what experimental conditions.

A striking empirical observation emerging from recent community-function studies is that microbial landscapes often appear surprisingly simple. In particular, linear regression analyses have shown that community-level functions can be predicted with high accuracy using only additive and pairwise terms, with higher-order contributions explaining little additional variance [9]. This apparent simplicity of small-size community-function landscapes has been reported across diverse experimental systems and functions, suggesting that higher-order interactions contribute little to the functional organization of many microbial communities.

Notably, similar patterns of apparent simplicity have also been reported in a seemingly distant domain: protein sequence–function landscapes. There, it has been argued that the apparent dominance of low-order structure may reflect not only underlying biological constraints, but also methodological choices, particularly regarding to how interaction effects are inferred from truncated models, and how experimental noise and global nonlinearities in the sequence–function mapping propagate into higher-order terms [19, 20]. However, others have argued that such nonlinear mappings may absorb or mask specific higher-order interactions, leading to an apparent but potentially misleading simplicity [21]. In high-dimensional landscapes, it is possible that the dominance of low-order epistatic terms may be a simple structural consequence of representation and sampling. Higher-order interactions might contribute little to the total variance simply because there are fewer of them in community-function landscapes of relatively small size. In turn, it is possible that measurement error and other sources of experimental noise impose an additional, order-dependent limit of detection.

Motivated by this hypothesis, we study a quantitative theoretical and empirical framework, borrowed from genetics, to characterize the structural and statistical limits of what can be inferred about ecological interactions from ecological landscapes. Armed with this framework, we ask which specific local (i.e. Taylor) interactions can be reliably detected at a given experimental noise level and replication depth, and we can also determine their functional relevance. Does the observed dominance of low-order interactions reflect a genuine concentration of functional signal at low orders, or does it arise from the combined effects of experimental noise and uncertainty propagation that masks higher-order contributions?

To address these two questions, we test the predictions of our framework across empirical and mechanistic settings. We first construct a large experimental community-function landscape comprising all 2^10^ possible communities of ten species (see Materials and Methods and SI, Section S2 for details). This scale allows us to directly assess how functional variance and detectability change with interaction order, and to identify the point at which biological signal becomes masked by the combinatorial amplification of noise under realistic experimental conditions.

Finally, to demonstrate the implications of our theoretical framework in a controlled setting, we analyze a synthetic landscape generated by the generalized Lotka-Volterra model. Because this model provides a fully deterministic, noise-free mapping from community composition to function with a known interaction structure, it serves as an ideal testbed to verify that, in the absence of measurement noise, our framework correctly recovers the expected low-order organization. We then add noise to this landscape to demonstrate how higher-order interaction signatures can arise spuriously from uncertainty propagation, even when the underlying dynamics contain only low-order interactions.

While the first question concerns notions of epistasis that are well established in genetics, revisiting them in ecology within a unified and minimal framework allows us to make explicit how noise, replication depth, and combinatorial structure constrain what can be inferred at each interaction order for microbial communities. Thus, we provide a unifying perspective bringing ideas from genetics to clarify ongoing discussions in microbial ecology.

## RESULTS

We represent community composition by a binary vector **x** ∈ {0, 1} ^*N*^, where *x*_*i*_ = 1 (*x*_*i*_ = 0) denotes the presence (absence) of species *i*. The community–function landscape *F* (**x**) assigns a quantitative function to each configuration. In experimental datasets, the biological signal 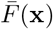 is not directly observable. Instead, as illustrated in Fig. 1C, each community configuration is typically measured a finite number of times, 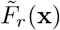, where *r* = 1, …, *R* indexes experimental replicates or batches, with *R* usually ranging from one to a few independent measurements per configuration. These observations are affected by stochastic variability and systematic batch-to-batch effects arising from experiments performed on different days. Throughout this work, we therefore analyze batch-corrected data, such that the residual fluctuations *ξ*_*r*_(**x**) represent stochastic variability around a common biological signal (see Fig. 1C and SI, Section S3),

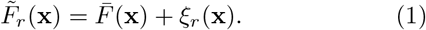

This decomposition assumes only that residual fluctuations have zero mean and finite variance; we allow for heteroscedasticity and examine the effect of correlated residuals explicitly in SI Section S7. Any such deviations from idealized noise assumptions can only increase uncertainty propagation and therefore reinforce our detectability limits. We now turn to the analysis of interaction structure in such noisy, high-dimensional landscapes.

### Detectability of local epistatic interactions is exponentially constrained by noise

Local epistatic interactions are obtained by combining function values across specific subsets of community configurations (see Fig. 1D), evaluated within the same experimental replicate *r*. For a given background configuration **B**, the epistatic interaction among a set *S* of *k* = |*S*| focal species—encoded by a binary vector **s** ∈ {0, 1} ^*N*^ with *s*_*i*_ = 1 for *i* ∈ *S*—is defined as an alternating sum over the 2^*k*^ community configurations obtained by toggling those species while keeping the background fixed (see Fig. 2A). This definition corresponds to a discrete, Taylor-like expansion of the landscape around the background **B**. Throughout, we implicitly identify subsets *U* ⊆ *S* with their corresponding indicator vectors **u** ∈ {0, 1} ^*N*^, so that **B** + **u** denotes the community configuration obtained by adding the species in *U* to the background. This yields

**FIG. 2.**
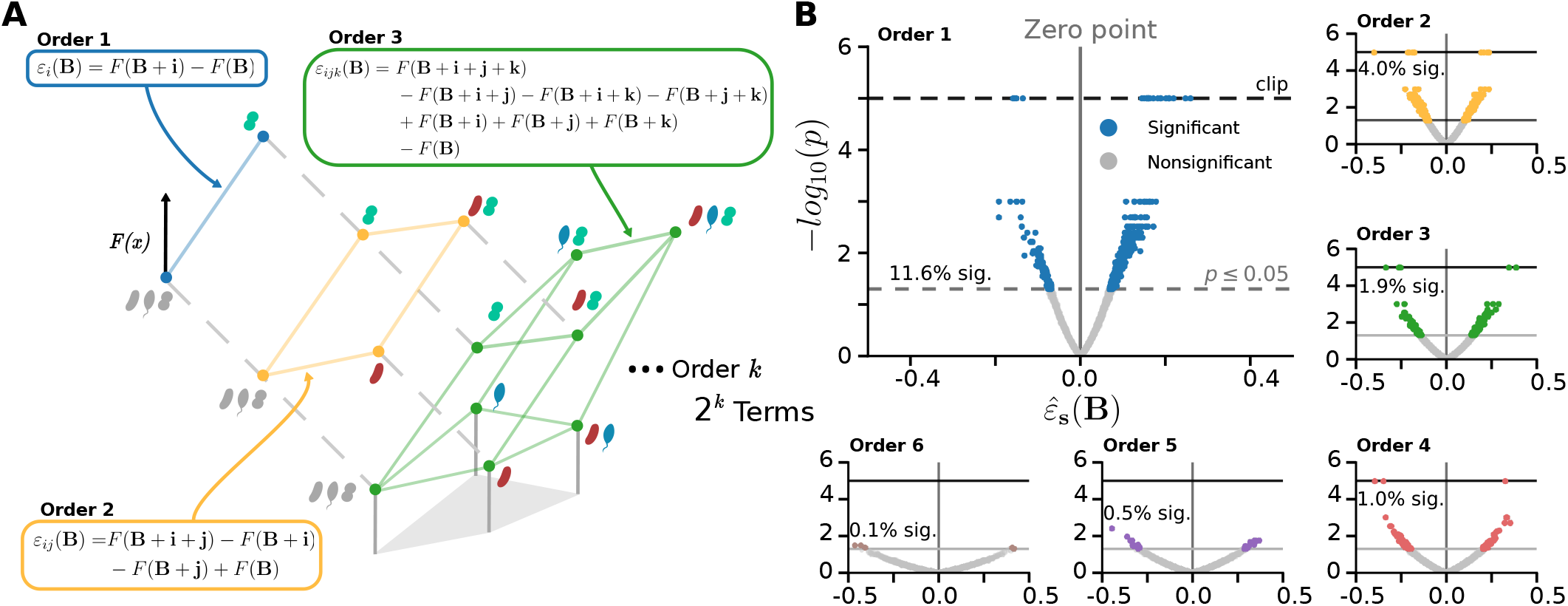
Detectability of local epistatic interactions is exponentially constrained by noise. **(A)** Schematic illustration of local epistatic interactions of increasing order evaluated in a fixed ecological background **B**. First-, second-, and third-order interactions are defined as alternating sums over 2^*k*^ community configurations obtained by toggling *k* focal species while keeping the background fixed. The number of terms entering the interaction grows exponentially with order, leading to combinatorial accumulation of experimental noise. **(B)** Volcano plots of local epistatic coefficients 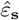(**B**) for interaction orders *k* = 1 to *k* = 6 computed from the fully sampled experimental community-function landscape described in the main text (*N* = 10 species, five biological replicates, see Materials and Methods and Section S2 of the SI). Orders *k* = 7–10 are omitted for clarity; the complete set of orders is shown in Fig. S6 of the SI. Each point corresponds to a specific set of focal species and background configuration. Although higher-order interactions span larger absolute magnitudes due to noise accumulation, their statistical significance rapidly deteriorates with increasing order. Dashed lines indicate the significance threshold (*p* ⩽ 0.05), and percentages denote the fraction of coefficients exceeding this threshold.

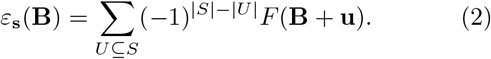

Each of the 2^*k*^ measurements contributes cumulatively to the dispersion of *ε*_**s**_(**B**), leading the associated variance to scale as ∼2^*k*^. This exponential amplification of uncertainty implies that, beyond a certain interaction order, real biological effects may become statistically indistinguishable from noise at the level of local interaction estimates.

To empirically investigate this issue, we constructed a fully-sampled community-function landscape by forming every possible combination of *N* = 10 species, growing all 1023 communities in minimal media (excluding the all-absent community **x** = **0**, which yields no growth/OD signal; see Materials and Methods), and determining their optical density as a function of choice. This landscape was constructed in five biological replicates, each of which represented a different experimental batch. We then used equations (1-2), as explained in the Materials and Methods and the SI, to quantify Taylor epistatic interactions at every order. The volcano plot representation in Fig. 2B contrasts the measured interaction strength with its statistical uncertainty. Statistical significance was assessed using a non-parametric, bootstrap-based null model. Under this null hypothesis, the biological signal is assumed to be absent, 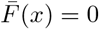, so that any apparent structure arises solely from experimental noise. Null landscapes are generated by resampling the batch-corrected residuals *ξ*_*r*_(*x*) in a way that preserves their empirical variance, heteroscedasticity, and any correlations across community configurations, while removing any systematic biological signal (see SI, Sections S5 and S7).

As interaction order increases, two clear effects emerge: individual epistatic coefficients tend to span a broader range of absolute values due to the accumulation of noise contributions, while the width of the corresponding null distribution grows even faster as a result of combinatorial amplification. Consequently, higher-order interactions may appear larger in magnitude yet become progressively harder to distinguish from noise. The majority of higher-order Taylor interactions are not significant, and this is even before correcting for multiple-hypothesis testing. Thus, we find that experimental noise precludes us from reaching conclusions about the strength of higher-order interactions in our experimental system.

### Why ecological landscapes appear simple: structural suppression of higher-order interactions

A recent theoretical analysis of various small-size community-function landscapes has found that additive and low-order interactions appear to dominate functional variance, while higher-order terms contribute little additional variance [9]. This pattern can be interpreted as evidence that biological interactions are intrinsically low-order. However, an alternative hypothesis is that the apparent simplicity of ecological landscapes arises naturally from the geometry of the combinatorial representation and from statistical constraints on inference, even in the presence of higher-order biological interactions.

Assessing whether higher-order interactions are important at the landscape level requires a representation that (i) averages over ecological contexts, (ii) assigns a unique contribution to each interaction order, and (iii) decomposes total functional variance in an orthogonal and non-redundant way. Such a representation naturally emerges when shifting to the Walsh–Hadamard (WH) basis, or equivalently to the Fourier transform of the landscape (see SI, Section S1) [22–24]. Each WH coefficient *f*_**s**_ corresponds to a subset **s** of focal species, and a WH coefficient of order *k* = |**s**| can be written as a uniform average of the corresponding local *k*-way interaction over all backgrounds [24],

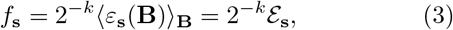

where *ε*_**s**_(**B**) denotes the local *k*-way epistatic interaction defined in Eq. (2) and ⟨·⟩_**B**_ denotes the uniform average over all background configurations. For later convenience, we defined *ℰ*_**s**_ = ⟨*ε*_**s**_(**B**)⟩_**B**_ as the background-averaged epistatic interaction associated within the focal species in **s**.

Because the WH functions form an orthogonal basis of the landscape (see Eq. (3)), the total functional variance admits a unique decomposition into contributions associated with different interaction orders [25, 26]. Specifically, the variance explained by all *k*-way interactions is defined as

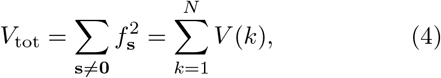

where 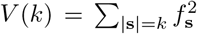 (Parseval’s theorem [27]). As shown in Fig. 3A, the empirical variance spectrum for our *N* = 10 landscape is strongly dominated by low-order interactions. First- and second-order terms account for most of the total functional variance, while contributions from higher orders rapidly decay and fall within the null expectation.

**FIG. 3.**
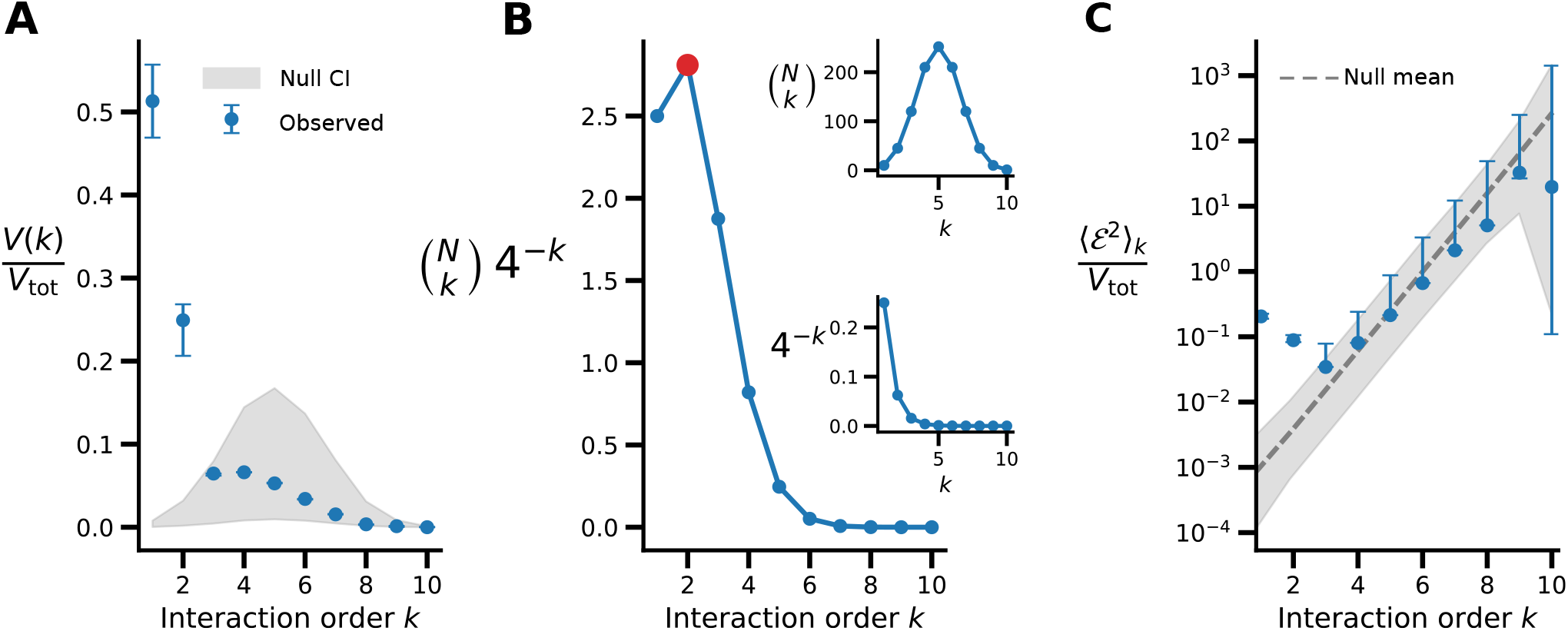
Limits of detectability and functional relevance across interaction orders. The figure provide a graphical decomposition of the order-resolved variance *V* (*k*) defined in Eq. (5), with panel **(A)** showing the variance spectrum itself and panels **(B)** and **(C)** displaying its two multiplicative components. **(A)** Order-resolved variance spectrum *V* (*k*)*/V*_tot_, showing the fraction of total functional variance explained by interactions of order *k*. Low-order interactions dominate the variance, while higher-order contributions fall within the 95% null confidence interval. **(B)** Structural factors controlling the variance spectrum. The combinatorial factor 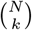 (top inset) grows rapidly with interaction order, whereas the geometric dilution factor 4^*−k*^ (bottom inset) suppresses high-order contributions. Their product, 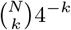, exhibits a maximum at *k* = 2 (highlighted in red) and decays thereafter, biasing the variance spectrum toward low-order interactions even if the epistatic amplitude of higher-order interactions truly increases. **(C)** Order-resolved epistasis amplitude ⟨ℰ^2^⟩_*k*_ normalized by total variance, compared to the 95% null expectation. Amplitudes for *k* = 1, 2 are clearly significant, whereas *k* = 3 lies on the border of the null model, suggesting a limit of detection. Crucially, a significant amplitude at order *k* indicates a detectable collective signal at that order, but does not guarantee that every individual interaction of that order can be resolved from noise. Qualitatively similar results for different ecological landscapes in the literature are illustrated in Fig. S9.

To understand the origin of this pattern, we rewrite the order-resolved variance using the relation between WH coefficients and background-averaged epistatic interactions Eq. (3) (see SI, Section S1),

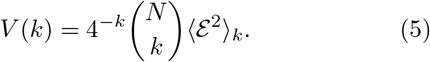

Here, ⟨ℰ^2^⟩_*k*_ denotes the order-*k epistasis amplitude*. Equation (5) shows that variance contributions are controlled by three factors: a combinatorial term 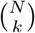 counting how many average-interactions exist at order *k*, a geometric prefactor 4^*−k*^ arising from the normalization of the WH coefficients, and the average strength of background-averaged *k*-way interactions captured by ⟨ℰ^2^⟩_*k*_. It is this latter term what we really care about, as it is the one which reflects the actual strength of biological interactions at that order.

The decomposition in Eq. (5) reveals a fundamental structural constraint. As illustrated in Fig. 3B, the product of the combinatorial and geometric dilution factors, 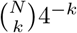, exhibits a broad maximum at *k* = 2 and decays thereafter, intrinsically biasing the variance spectrum toward low-order interactions. This bias arises independently of the underlying biological interaction structure, and it is simply dependent on the size of the community-function landscape. It is straightforward to see that, as the size of the community-function landscape increases beyond N=10, the maximum of 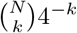 will trivially shift to higher and higher interaction orders *k*, simply because there are a growing number of these interactions.

The remaining contribution, the epistasis amplitude ⟨ℰ^2^⟩_*k*_ shown in Fig. 3C, appears to increase with interaction order for our empirical landscape. However, this trend should not be interpreted as evidence for intrinsically stronger higher-order biological interactions. In fully sampled landscapes, background averaging ensures that the estimation uncertainty of individual Walsh–Hadamard coefficients *f*_**s**_ is approximately independent of *k*. Because background-averaged epistatic interactions scale as ℰ_**s**_ = 2^*k*^*f*_**s**_ (Eq. (3)), any residual estimation error is exponentially amplified with interaction order.

As a result, noise alone is sufficient to generate an apparent growth of epistasis amplitudes at high orders. Consistently, only first- and second-order interactions are clearly distinguishable from the null model (Fig. 3C). At third order, the epistasis amplitude lies at the boundary of the null distribution, while higher-order contributions are fully indistinguishable from noise. This noise-driven amplification obscures the true behavior of underlying biological interactions and fundamentally limits reliable inference of high-order interactions in community-function landscapes.

It is important to remark that even if the observed 4^*k*^ growth of epistasis amplitudes with interaction order in Fig. 3C were not an artifact, and it did in fact reflect genuine biological interactions, still the contribution of high-order interactions to the total functional variance would be small, as it would be suppressed by the structural prefactor. Therefore, even in small community-function landscapes that contain large high-order interactions, the variance of the landscape will still be dominated by first- and second-order interactions, while higher-order terms remain functionally negligible.

Taken together, Fig. 3 show that the dominance of low-order interactions arises from the combined effects of combinatorial scaling, geometric dilution, and noise amplification. In principle, increasing the number of species *N* would shift the maximum of the structural factor 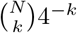 toward higher interaction orders. In practice, however, a fully sampled ecological landscape with *N* = 10 species already lies close to the experimental limit of what can be feasibly constructed and replicated in the laboratory. Moreover, even if substantially larger landscapes could be constructed, the statistical constraints identified here would still prevent reliable inference of higher-order biological interactions. The same qualitative behavior of the variance spectrum and epistasis amplitudes is observed across several additional microbial community-function landscapes analyzed in previous studies [9] (see SI Section S6 and Fig. S9), indicating that this pattern is not unique to our empirical landscape (it must be noted that the arguments we have presented hold for fully and uniformly sampled landscapes over {0, 1}^*N*^).

### Apparent higher-order interactions can arise from noise in pairwise systems

To disentangle the effects of experimental noise from those of the underlying ecological mechanism, we analyzed community–function landscapes generated by the generalized Lotka–Volterra (gLV) model (Fig. 4A). By construction, gLV dynamics include only pairwise ecological interactions: species interact exclusively through a bilinear interaction matrix, with no explicit higher-order terms in the governing equations (Fig. 4B). However, it is still possible that indirect effects might lead to the existence of non-zero higher-order epistatic interactions (i.e. non-zero Walsh-Hadamard coefficients for *k >* 2). Here we are distinguishing explicitly between higher-order interactions in the ecological dynamics (defined as explicit multi-species terms in the equations of motion) and higher-order structure in the resulting community– function landscape.

**FIG. 4.**
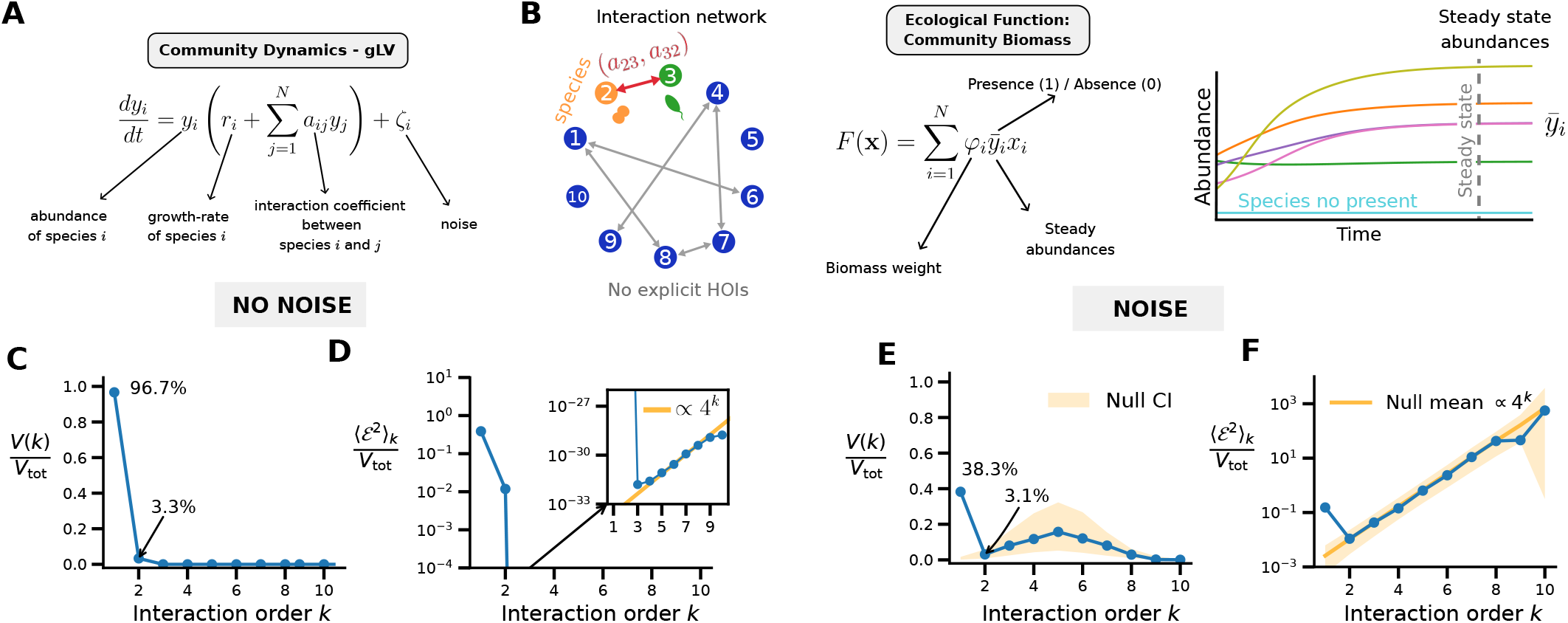
Noise can generate apparent higher-order structure in a purely pairwise ecological model. **(A)** Generalized Lotka–Volterra (gLV) dynamics used to generate synthetic community–function landscapes. Species interact exclusively through pairwise terms *a*_*ij*_; no explicit higher-order interaction terms are present in the dynamics. **(B)** Representative interaction network illustrating the pairwise structure of the model. The community function is defined as total biomass at steady state, 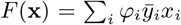, where *x*_*i*_ ∈ {0, 1} denotes de presence/absence of species *i* in the dynamics, 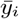 denotes steady-state abundances and *φ*_*i*_ is the biomass weight. **(C)** Order-resolved variance spectrum *V* (*k*)*/V*_tot_ of the noise-free landscape, showing that functional variance is almost entirely explained by first- and second-order terms. **(D)** Corresponding epistasis amplitude, confirming the absence of intrinsic higher-order structure. **(E)** Variance spectrum after adding experimental noise, revealing a broad peak at intermediate orders despite the absence of genuine higher-order interactions. Shaded regions indicate the 95% confidence interval of a noise-only bootstrap null model. **(F)** Epistasis amplitude after adding experimental noise exhibits a characteristic 4^*k*^ scaling arising from noise propagation and lies entirely within the 95% null expectation, indicating that the observed higher-order signal is consistent with measurement noise alone.

As hinted above, higher-order epistatic interactions in the community-function landscape could still arise indirectly, even in a purely pairwise dynamical system, because steady states involve the inversion of the interaction matrix [28]. However, in the absence of experimental noise, such effects are negligible across the parameter regime explored here (Fig. 4C,D). The resulting landscapes are intrinsically low order: nearly all functional variance is captured by first- and second-order interactions, with higher-order contributions being numerically negligible (Fig. 4C,D). Thus, a purely pairwise ecological mechanism produces a genuinely low-dimensional functional landscape.

We then modeled the presence of “experimental noise” by perturbing steady-state abundances and averaging the resulting function over a finite number of replicates. In this noisy regime, the observed variance spectrum changes quantitatively. While the underlying ecological dynamics remain strictly pairwise, the higher-order epistatic terms reflecting this noise become increasingly prominent, generating a broad peak at intermediate interaction orders (Fig. 4E,F).

Together, these results provide a mechanistic demonstration that experimental noise alone can generate the illusion of dominant higher-order interactions in community–function landscapes. Apparent high-order structure in empirical data should therefore not be necessarily interpreted as evidence of hidden ecological complexity, but rather as a consequence of noise amplification and limited detectability in high-dimensional composition spaces.

## DISCUSSION

Inspired by fitness landscapes in genetics, highdimensional community–function landscapes offer a natural framework to describe how collective ecological functions emerge from composition. Here, we showed that the inference of epistatic interactions in such landscapes is constrained by two distinct and unavoidable effects. Just as it has been reported in genetics, for local, background-dependent interactions uncertainty grows exponentially with interaction order because each *k*-way interaction is built from an alternating combination of 2^*k*^ community configurations, leading to rapid noise accumulation (Fig. 2). However, when interactions are assessed through their functional contribution, the dominance of low orders in the variance spectrum emerges naturally from the interplay of combinatorial multiplicity and geometric dilution (Eq. (5); Fig. 3), with experimental noise further obscuring high-order structure.

Recent work on protein sequence–function landscapes has highlighted an active debate on the interpretation of apparent low-order structure, with contrasting views on whether higher-order epistasis is genuinely weak or instead masked by methodological choices such as reference selection and nonlinear mappings [19–21]. Our results place this debate in a broader and more general framework. Rather than attributing apparent simplicity to specific modeling choices, we show that uncertainty propagation and signal suppression are structural consequences of how interactions are constructed and measured in high-dimensional landscapes.

Distinguishing genuine biological signal from noise has direct implications for applied microbial ecology and biotechnology. Our results place recent empirical observations—that microbial community–function landscapes are dominated by additive and pairwise structure and are highly compressible [9, 29]—on firm statistical and mechanistic grounds. From a biotechnological perspective, this implies that engineering strategies relying on higher-order interaction complexity are not only experimentally costly, but also statistically unreliable under realistic sampling regimes, as contributions with *k* ⩾ 3 are typically indistinguishable from noise. In contrast, additive and pairwise interactions (*k* = 1, 2) concentrate both detectable signal and explainable variance, providing a principled and practically actionable foundation for rational community design. Notably, an analogous conclusion has long been recognized in genetic fitness landscapes, where low-order epistasis is sufficient to constrain evolutionary trajectories and global landscape structure despite the combinatorial explosion of possible interactions [30, 31].

Although our analysis is framed in terms of ecological landscapes, the constraints we identify arise from the generic structure of high-dimensional composition– to–function mappings and therefore extend well beyond microbial ecology. Similar questions regarding the relevance, scaling, and detectability of higher-order epistasis arise across a wide range of systems, including protein sequence–function landscapes [32], genotype–phenotype maps in protein and gene-regulatory models [33], tRNA fitness landscapes [34], the genetic architecture of complex traits [35], and evolutionary trajectories in systems such as yeast and bacteriophages [36].

Ultimately, we make no claim about the absence of higher-order interactions in nature. Rather, we show that their existence can be challenging to established from finite, noisy measurements like the ones allowed by modern technology for community-function landscapes. Even when higher-order effects are present in the underlying biology, the combinatorial structure of interaction estimators and the propagation of measurement uncertainty impose intrinsic, order-dependent limits on detectability. As a result, only a restricted level of interaction complexity can be rigorously inferred from realistic experimental datasets.

Lastly, we note that our analysis is formulated for experimentally constructed community–function landscapes, where compositions are defined by presence/absence in controlled assembly assays. Extrapolating to natural microbial communities requires caution, given natural habitats add spatial structure, environmental heterogeneity, and context-dependent dispersal that are not represented in our framework. It is possible, however, that many of these additional ecological constraints may further bias effective interactions toward low orders. In spatially structured environments such as soils and biofilms, interactions are often mediated by diffusion and local co-localization, which can limit the number of species that effectively couple within the same microenvironment [37, 38], driving mechanistic descriptions of microbial communities to a regime akin to our Lotka-Volterra scenario. While such constraints do not eliminate higher-order effects in principle, they may reduce their effective strength and prevalence, acting in the same direction as the detectability limits quantified here and reinforcing the dominance of additive and pairwise contributions.

## MATERIALS AND METHODS

### Software availability

All analyses were performed using the Epistasia Python package, developed by the authors and available at https://github.com/MCMateu/Epistasia. Documentation is provided at https://mcmateu.github.io/Epistasia/.

### Experimental design and community assembly

Ten bacterial strains were a kind gift by Ramon Santamaria and Beatriz Santos, were originally isolated from a natural beehive community. We assembled all possible combinations of *N* = 10 species using the assembly methods and protocols developed in [11]. All species were inoculated at a starting OD_600_ of 0.01, and the communities were cultured for 48h at 28 °C in 500 *µL* LB medium inside 96-well deep-well plates. Community function was quantified for all communities at the same time as total optical density through 200 *µL* in a separate 96 well plate, after homogeneization (OD_600_). The experiments were conducted in *R* = 5 biological replicates in as many batches, on separate days and corrected for batch effect. Full experimental details are provided in the SI, Section S2.

## ACKNOWLEDGEMENTS

This work has been funded by the European Union (ERC, ECOPROSPECTOR, 101088469) and Juan de la Cierva grant JDC2024-054148-I. The views and opinions expressed here are, however, those of the author(s) only and do not necessarily reflect those of the European Union or the European Research Council. Neither the European Union nor the granting authority can be held responsible for them. We thank Ramon Santamaria and Beatriz Santos for donating the strains we used to construct the community-function landscape, and we also thank Juan Díaz-Colunga and Alfonso Mendaña for critical reading of the manuscript, Andrea Arrabal and Belén Benítez-Domínguez for scientific discussions, and Alba Rodriguez Carreño for logistic and administrative assistance at all stages of this project. We would like to dedicate this manuscript to the memory of Professor Kevin Wood, whose pioneering work on epistatic-like interactions outside the realm of fitness landscapes inspired the development of community-function landscapes and all the subsequent work we have conducted on this field.

