## Supplementary Information for "On the limits of detection of epistatic higher-order interactions"

#### Contents

S1. Definitions and notation 2
- S1.1. Community function landscapes 2
- S1.2. Interactions and epistasis 2
- S1.3. Walsh-Hadamard basis 2
- S1.4. Variance decomposition across interaction orders 3
- S1.5. Walsh–Hadamard expansion under incomplete sampling 3

S2. Experimental methods 4

S3. Measurement noise and batch effects 6
- S3.1. Measurement model 6
- S3.2. Batch effects removal 6

S4. Propagation of measurement noise to interaction measures 7
- S4.1. Epistatic interactions 7
- S4.2. Noise projection on the Walsh–Hadamard coefficients 8
- S4.3. Contribution of noise to order-specific variances 8

S5. Uncertainty quantification and limits of detectability 8
- S5.1. Bootstrap-based uncertainty estimation 9
- S5.2. Bootstrap null model 9
- S5.3. Bootstrap $p$-values for a generic statistic $Q(\mathbf{x})$ 9
- S5.4. Detectability limits for higher-order interactions 10

S6. Analysis of previously published datasets 10

S7. Extension to correlated residuals 10

S8. Synthetic community landscapes 12
- S8.1. Random maps 12
- S8.2. Flat epistatic landscapes 13
- S8.3. Mechanistic testbed: generalized Lotka–Volterra landscapes 13

References 14

S9. Supplementary Tables 15

S10. Supplementary Figures 15

### S1 Definitions and notation

##### S1.1. Community function landscapes

We consider ecological systems composed of up to $N$ species, and describe each community configuration by a binary vector $\mathbf{x} = (x_1, \ldots, x_N)$, where $x_i = 1$ or $0$ indicates the presence or absence of species $i$. The set of all possible communities thus forms the discrete space $\{0,1\}^N$, containing $2^N$ distinct configurations. To each configuration we associate a quantitative community-level outcome, such as total biomass, growth rate, or ecosystem productivity. This defines a *community-function landscape*, a mapping $F : \{0,1\}^N \to \mathbb{R}$ that assigns a functional value to every possible combination of species. Experimentally, such landscapes are sampled by measuring the function of multiple communities, often with $R$ replicated measurements for the same configuration $\mathbf{x}$.

##### S1.2. Interactions and epistasis

The effect of a species on a community-level function generally depends on the presence or absence of other species. This context dependence is formalized by defining interactions relative to a fixed *background* $\mathbf{B}$, representing the configuration of all species except a chosen set of *focal* ones. Mathematically, for a set of focal species $\mathcal{A}$, the background $\mathbf{B}$ is a binary vector such that all focal species are absent, $B_i = 0$ for all $i \in \mathcal{A}$. The simplest case corresponds to a single focal species, $\mathcal{A} = \{i\}$. For a given background $\mathbf{B}$, the first-order effect of species $i$ is defined as the finite difference

$$\Delta_i F(\mathbf{B}) = F(\mathbf{B} + \mathbf{i}) - F(\mathbf{B}), \tag{1}$$

where $\mathbf{i}$ denotes the unit vector with a 1 at position $i$ and zero otherwise. This measures the marginal contribution of adding species $i$ to the community $\mathbf{B}$. Second-order (pairwise) interactions quantify the non-additive combined effect of two species on the community-level function. Thus, pairwise epistasis between species $\mathcal{A} = \{i, j\}$ in background $\mathbf{B}$ (where $B_i = B_j = 0$) is defined as

$$\varepsilon_{ij}(\mathbf{B}) = F(\mathbf{B} + \mathbf{i} + \mathbf{j}) - F(\mathbf{B} + \mathbf{i}) - F(\mathbf{B} + \mathbf{j}) + F(\mathbf{B}), \tag{2}$$

which corresponds to the standard $F_{11} - F_{10} - F_{01} + F_{00}$ definition widely used in the epistasis literature, with the background made explicit [1]. More generally, a *kth-order interaction* captures an irreducible collective effect that involves exactly $k$ species and cannot be decomposed into contributions from interactions of lower order. Let $S \subset \{1, \ldots, N\}$ be a subset of $k = |S|$ focal species, encoded by a binary indicator vector $\mathbf{s} \in \{0,1\}^N$ with $s_i = 1$ for $i \in S$. We consider a background configuration $\mathbf{B}$ in which all focal species are absent, i.e. $B_i = 0$ for all $i \in S$. We implicitly identify any subset $U \subseteq S$ with its corresponding indicator vector $\mathbf{u} \in \{0,1\}^N$, where $u_i = 1$ if $i \in U$ and $u_i = 0$ otherwise. The corresponding $k$-way epistatic interaction in background $\mathbf{B}$ is then defined as the alternating sum over the $2^k$ community configurations obtained by toggling the focal species,

$$\varepsilon_{\mathbf{s}}(\mathbf{B}) = \sum_{U \subseteq S} (-1)^{k-|U|} F(\mathbf{B} + \mathbf{u}). \tag{3}$$

where the sum runs over all subsets $U \subseteq S$, and $\mathbf{B} + \mathbf{u}$ denotes the community configuration obtained by adding to the background $\mathbf{B}$ all species in $\mathbf{u}$ (i.e. setting $x_i = 1$ for $i \in U$ while leaving all other components unchanged). Here, $F(\mathbf{x})$ denotes the community–function value associated with configuration $\mathbf{x}$. By construction $\varepsilon_{\mathbf{s}}(\mathbf{B})$ vanishes whenever the contribution of these species can be fully expressed in terms of interactions of order strictly smaller than $k$.

##### S1.3. Walsh-Hadamard basis

To analyze interactions across different orders in a unified and systematic way, we represent the community-function landscape in the Walsh–Hadamard (WH) basis, which provides an orthogonal decomposition for functions defined on the binary space $\{0,1\}^N$. In this representation, each basis function is naturally associated with a specific subset of species, and therefore with a well-defined interaction order. We work with the shifted WH basis functions

$$\phi_{\mathbf{s}}(\mathbf{x}) = (-1)^{\mathbf{s} \cdot (\mathbf{x} + \mathbf{1})}, \tag{4}$$

where $\mathbf{s} \in \{0,1\}^N$ labels the subset of species involved. Any landscape $F(\mathbf{x})$ can be uniquely expanded as

$$F(\mathbf{x}) = \sum_{\mathbf{s}} f_{\mathbf{s}} \, \phi_{\mathbf{s}}(\mathbf{x}), \tag{5}$$

with coefficients

$$f_{\mathbf{s}} = 2^{-N} \langle \phi_{\mathbf{s}} | F \rangle, \tag{6}$$

where $\langle\cdot|\cdot\rangle$ denotes the usual inner product over $\{0,1\}^N$, $\langle g|f\rangle = \sum_{\mathbf{x}} g(\mathbf{x})f(\mathbf{x})$. Crucially, the WH coefficients admit a direct ecological interpretation. The constant mode $f_{\mathbf{0}}$ corresponds to the mean function across all community configurations. Modes involving a single species capture its average additive effect across all backgrounds. More generally, a mode involving $k$ species corresponds to the average $k$-way epistatic interaction among those species, evaluated over all possible backgrounds of the remaining community. Formally, the coefficient associated with a subset $\mathbf{s}$ of size $|\mathbf{s}| = k$ is proportional to the background-averaged local epistasis,

$$f_{\mathbf{s}} = 2^{-k}\left(\frac{1}{2^{N-k}}\sum_{\mathbf{B}\in\{0,1\}^{N-k}} \varepsilon_{\mathbf{s}}(\mathbf{B})\right) = 2^{-k}\langle\varepsilon_{\mathbf{s}}(\mathbf{B})\rangle_{\mathbf{B}}, \tag{7}$$

where $\varepsilon_{\mathbf{s}}$ denotes the local $k$-way epistatic interaction defined in the previous section, and $\langle\cdot\rangle_{\mathbf{B}}$ denotes the uniform average over all background configurations $\mathbf{B}$. We will therefore refer to $\mathcal{E}_{\mathbf{s}} \equiv \langle\varepsilon_{\mathbf{s}}(\mathbf{B})\rangle_{\mathbf{B}}$ as the *interaction strength* associated with subset $\mathbf{s}$.

##### S1.4. Variance decomposition across interaction orders

The variance decomposition presented below is exact only under the assumption that the full configuration space $\{0,1\}^N$ is sampled uniformly. The WH expansion provides a natural decomposition of the total functional variance of a landscape across interaction orders. Let

$$V_{\text{tot}} = \left\langle (F - \langle F\rangle)^2 \right\rangle_{\mathbf{x}} \tag{8}$$

denote the total variance of the community-function landscape, where $\langle\cdot\rangle_{\mathbf{x}}$ denotes the uniform average over all community configurations $\mathbf{x} \in \{0,1\}^N$. By orthogonality of the WH basis, this variance can be written as a sum of squared coefficients,

$$V_{\text{tot}} = \sum_{\mathbf{s}\neq\mathbf{0}} f_{\mathbf{s}}^2 = \sum_{k=1}^{N} V(k), \tag{9}$$

where the contribution of interaction order $k \geq 1$ is

$$V(k) = \sum_{\substack{\mathbf{s}\in\{0,1\}^N:\\ |\mathbf{s}|=k}} f_{\mathbf{s}}^2. \tag{10}$$

Using the relation between Walsh–Hadamard coefficients and background-averaged epistatic interactions Eq. (7), this quantity can be written as

$$V(k) = 4^{-k}\sum_{\substack{\mathbf{s}:\\ |\mathbf{s}|=k}} \mathcal{E}_{\mathbf{s}}^2 = 4^{-k}\binom{N}{k}\langle\mathcal{E}^2\rangle_k, \tag{11}$$

where

$$\langle\mathcal{E}^2\rangle_k = \binom{N}{k}^{-1} \sum_{\substack{\mathbf{s}\in\{0,1\}^N:\\ |\mathbf{s}|=k}} \mathcal{E}_{\mathbf{s}}^2 \tag{12}$$

defines the *order-$k$ epistasis amplitude*. This quantity measures the typical strength of $k$-way interactions, independently of how many such interactions exist. The variance contribution $V(k)$ results from the product of this intrinsic amplitude and the combinatorial factor $\binom{N}{k}$, which counts the number of possible $k$-way interactions. As a consequence, higher-order interactions can in principle contribute substantially to the total variance even if their typical strength is modest. The variance spectrum $\{V(k)\}_k$ thus provides a compact summary of how functional variability is distributed across interaction orders, as illustrated in Fig. 3 of the main text.

##### S1.5. Walsh–Hadamard expansion under incomplete sampling

The orthogonality of the WH basis and the associated variance decomposition rely on the landscape being defined on the full hypercube $\{0,1\}^N$ under the uniform measure. We now consider the case in which the landscape is observed only on a subset of configurations. Let $\Omega \subsetneq \{0,1\}^N$ denote the set of observed community configurations. In this case, the natural inner product is defined only over the observed subset,

$$\langle F, G\rangle_\Omega = \sum_{\mathbf{x}\in\Omega} F(\mathbf{x})G(\mathbf{x}), \tag{13}$$

which corresponds to the uniform empirical measure over $\Omega$. Under this restricted inner product, the WH basis functions $\phi_{\mathbf{s}}$ are no longer orthogonal. The corresponding Gram matrix is

$$G_{\mathbf{st}} = \langle \phi_{\mathbf{s}}, \phi_{\mathbf{t}} \rangle_\Omega = \sum_{\mathbf{x}\in\Omega} \phi_{\mathbf{s}}(\mathbf{x})\phi_{\mathbf{t}}(\mathbf{x}), \tag{14}$$

and in general $G_{\mathbf{st}} \neq \delta_{\mathbf{st}}$. Let $\mathbf{\Phi}$ denote the design matrix with entries $\Phi_{\mathbf{xs}} = \phi_{\mathbf{s}}(\mathbf{x})$ for $\mathbf{x} \in \Omega$. Then the Gram matrix can be written compactly as

$$\mathbf{G} = \mathbf{\Phi}^\top \mathbf{\Phi}. \tag{15}$$

Because the basis is not orthogonal, the variance no longer decomposes as a simple sum of squared coefficients. Instead,

$$\mathrm{Var}_\Omega(F) = \frac{1}{|\Omega|}\mathbf{f}^\top \mathbf{G}\mathbf{f} - \left(\frac{1}{|\Omega|}\mathbf{f}^\top \boldsymbol{\mu}\right)^2, \tag{16}$$

where $\mu_{\mathbf{s}} = \langle \phi_{\mathbf{s}}, 1\rangle_\Omega$ and the WH coefficients $\mathbf{f}$ are obtained by solving the normal equations

$$\mathbf{G}\mathbf{f} = \mathbf{\Phi}^\top \mathbf{F}. \tag{17}$$

Cross-terms encoded in $\mathbf{G}$ couple different interaction orders, and variance contributions cannot be uniquely assigned to a single order $k$. In particular, the quantities $V(k) = \sum_{|\mathbf{s}|=k} f_{\mathbf{s}}^2$ can still be computed, but they no longer represent orthogonal contributions to the variance. Rather, they are simply the sum of squared WH coefficients at order $k$ computed under the restricted support $\Omega$, without a direct geometric interpretation.

Orthogonality could in principle be restored by constructing a new basis that diagonalizes the Gram matrix $\mathbf{G}$ (e.g. via Gram–Schmidt orthogonalization or spectral decomposition). However, the resulting orthogonal modes would be linear combinations of WH functions and would no longer correspond to specific subsets of species. As a consequence, they would lose their direct ecological interpretation as $k$-way interactions. One may still define an empirical order-$k$ epistasis amplitude as

$$\langle \mathcal{E}^2 \rangle_{k,\Omega} = \binom{N}{k}^{-1} \sum_{\substack{\mathbf{s}\in\{0,1\}^N: \\ |\mathbf{s}|=k}} \mathcal{E}^2_{\mathbf{s},\Omega}, \tag{18}$$

where the average is taken over backgrounds for which all required configurations entering Eq. (3) are observed. However, unlike in the fully sampled case, $\langle \mathcal{E}^2 \rangle_{k,\Omega}$ is not an intrinsic geometric property of the combinatorial landscape but an empirical statistic. Because orthogonality is lost and the inner product depends on the sampling support $\Omega$, the amplitude becomes support-dependent. It therefore reflects both the underlying biological structure and the pattern of sampling, and is not directly comparable across datasets with different coverage.

#### S2 Experimental methods

##### Community origin and strain isolation

The starting material for this study consisted of environmental bacterial communities associated with a single beehive located in the province of Salamanca (Spain). Samples were originally collected and processed by Dr. Ramón Santamaría at the Instituto de Biología Funcional y Genómica (IBFG, Salamanca). Briefly, material from the beehive was suspended and plated on a general-purpose complex medium (hereafter "Sharlau medium" (Nutrient broth, Sharlau, REF: 02-140-500)). For isolation, samples were plated on Scharlau medium solidified with $2\%$ (w/v) bacteriological agar and incubated at 28 ºC for 48 h. Under these conditions, colonies displayed a wide range of colours and morphologies. Colonies with distinct pigmentation and morphology were picked and re streaked on fresh Scharlau agar plates. This process was repeated at least twice to obtain purified isolates. From the resulting culture collection, ten isolates exhibiting the most clearly distinguishable colony colours and morphologies were selected for further characterization and for subsequent landscape experiments. These isolates are referred to as C1, C4, C5, C6, C7, C8, C9, C10, C12 and C14 throughout the manuscript. To ensure purity, each isolate was streaked for single colonies and passaged at least three times on Scharlau agar before being used in sequencing or community assembly experiments. Glylcerol stocks of isolates were prepared by mixing equal volumes of culture and $40\%$ glycerol and storing at $-80$ºC.

##### DNA sequencing and taxonomic assignment

To confirm the taxonomic identity of the isolates and to assess genetic differences among them, we sequenced the terminal regions of the 16S rRNA gene of each strain by Sanger sequencing. Briefly, a single, well-isolated colony grown on Scharlau agar was inoculated into 20 mL of LB medium (Luria Broth [Condalab, REF: 1551.00]) in a 50 mL Falcom tube and incubated overnight at 28 ºC and 200 rpm. 1mL of the overnigth culture was transferred to a 1.5 mL microcentrifuge tube and cells were harvested by centrifugation (25 min at 3,000 × g, room temperature), the supernatant was discarded, and the cell pellet was shipped on ice to the sequencing facility at Hospital Universitario Ramón y Cajal (Madrid, Spain).

At the facility, genomic DNA was extracted and two Sanger reactions were performed per isolate, each using a primer targeting one of the termini of the 16S rRNA gene. This yielded high-quality reads for the $5'$ and $3'$ ends of the 16S gene, whereas the central region between the two reads was only partially covered and therefore not suitable for reliable full-length consensus reconstruction. Chromatograms were imported into Geneious (Biomatters Ltd.) for processing. Forward and reverse reads were quality-checked and trimmed based on Phred scores, and only the high-quality segments at each end were retained for downstream analyses. Taxonomic assignment was performed by comparing these curated terminal 16S segments against the SILVA rRNA reference database using BLASTn. Species-level assignments for the ten isolates are summarized in Table S1 and were used as the basis for defining strains in subsequent analyses.

#### Sanger sequence processing and genetic distance analysis

Raw Sanger reads were processed in Geneious to obtain high-quality partial 16S rRNA sequences suitable for comparative analyses. Each chromatogram was trimmed according to its base-quality profile using Geneious' built-in quality trimming (modified Mott algorithm), and bases in low-quality central regions were discarded. When possible, overlapping portions of the two terminal reads were locally aligned to refine base calls in the overlap; only positions within these high-confidence terminal regions were retained for distance calculations.

To quantify genetic differences among isolates, the curated terminal 16S segments from both ends were aligned jointly in Geneious, and a global alignment was computed over the combined high-quality regions. For each pair of isolates, we counted the number of mismatched base pairs across all aligned high-confidence positions, yielding a symmetric matrix of pairwise SNP distances (reported in Table S2). Although these distances do not correspond to full-length 16S divergence, they are sufficient for species-level identification and for relative comparison among isolates.

Based on this matrix, isolates sharing the same species-level assignment but differing by more than 3 SNPs across the aligned terminal regions were operationally defined as distinct strains. Under this criterion, C1 and C12 (both assigned to Dermacoccus nishinomiyaensis) and C8 and C9 (both assigned to Rhodococcus corynebacterioides) were considered distinct strains and retained as separate entities in the synthetic landscapes.

#### Culture media and growth conditions

All landscape assembly and functional assays were performed in Scharlau medium (liquid formulation matching the solid medium used for isolation). For preculture, each strain was streaked from frozen glycerol stocks onto Scharlau agar plates ($2\%$ w/v agar) and incubated at 28 ºC for 48 h. Single colonies from these plates were then used to inoculate $20 - 25$ mL of liquid Scharlau medium in sterile culture tubes. Liquid cultures were grown overnight at 28 ºC with orbital shaking (e.g. 200 rpm). Optical density at 600 nm of overnight cultures ($OD_{600}$) was measured in a 96-well plate spectrophotometer using sterile Scharlau medium as blank, and the cell suspension was adjusted to an $OD_{600}$ of $0.1$. These OD standardized suspensions served as inocula for all landscape experiments.

#### Construction of 10 strain landscapes

For the construction of 10 strains landscape, five different identical replicates of a 8 species landscape are needed.

##### *Construction of backbone landscapes of 8 species*

Synthetic microbial landscapes containing 8 strains were generated by combinatorial mixing of the standardized inocula in 96 well deep well plates, following the protocol described in [2] with modifications tailored to this strain set and medium. For each 8 strain landscape, the final culture volume per well was 400 µL, with a minimum contribution of 50 µL from any strain present in a given well. The 8 species landscape was replicated five times by the same user in the same laboratory, on different days and starting from different colonies.

##### *Extension to 10 strain landscapes*

To construct landscapes including all 10 strains, the 8 strain assembled as described above were used as backbones, and the remaining two strains (hereafter referred to as species 9 and species 10; corresponding to isolates C12 and C14 ) were added in defined combinations:

- $L_1$ (backbone only): 100 $\mu$L of Scharlau medium were added to the 8 strain backbone mixture to reach a final volume of 500 $\mu$L .

- $L_2$ (backbone + species 9): 50 $\mu$L of the OD standardized inoculum of species 9 and 50 µL of Scharlau medium were added to the backbone.
- $L_3$ (backbone + species 9 + species 10): 50 $\mu$L of species 9 and 50 $\mu$L of species 10 were added to the backbone.
- $L_4$ (backbone + species 10): 50 $\mu$L of species 10 and 50 $\mu$L of Scharlau medium were added to the backbone.

All plates were incubated at 28 ºC for 48 h.

###### Measurement of community function

Community function was quantified as total biomass accumulation, using optical density at 600 nm ($OD_{600}$) as a proxy. After 48 h of incubation, cultures in deep well plates were homogenized by gently pipetting up and down several times with a multichannel pipette, taking care to resuspend the biomas (taking care to avoid the formation of bubbles). From each well, 150 $\mu$L of the resuspended culture were transferred to a sterile, flat bottom 96 well microtiter plate. $OD_{600}$ was measured using a microplate spectrophotometer with Scharlau medium as blank.

#### S3 Measurement noise and batch effects

##### S3.1. Measurement model

Experimental measurements of community-function landscapes are affected by stochastic noise and systematic batch-to-batch variability. We model the observed function $\tilde{F}_r(\mathbf{x})$ measured in replicate (or batch) $r$ as

$$\tilde{F}_r(\mathbf{x}) = \bar{F}(\mathbf{x}) + a_r + b_r\, \pi_r(\mathbf{x}), \tag{19}$$

where $\bar{F}(\mathbf{x})$ denotes the underlying biological signal –independent of the replicate $r$– associated with configuration $\mathbf{x}$. The parameters $a_r$ and $b_r$ capture additive and multiplicative batch effects, accounting for systematic shifts in mean and variance across experimental runs. The term $\pi_r(\mathbf{x})$ represents stochastic measurement noise –such as fluctuations in inoculum size or readout variability in biomass or optical density measurements–assumed to have zero mean and finite variance.

An example of the batch effects present in the empirical datasets analyzed in this work, and their removal using the procedure described below, is shown in Fig. 1. Before correction, experimental replicates exhibit systematic shifts in both mean and variance across batches. After batch correction, replicate distributions collapse onto a common scale, consistent with a shared underlying biological signal.

##### S3.2. Batch effects removal

We perform batch correction within a Bayesian generative framework defined directly on the measurement scale $\tilde{F}_r(\mathbf{x})$. Weakly informative priors are placed on the additive and multiplicative batch effects and on the stochastic noise,

$$a_r \sim \mathcal{N}(0, \sigma_a^2), \qquad \log b_r \sim \mathcal{N}(0, \sigma_b^2), \qquad \pi_r(\mathbf{x}) \sim \mathcal{N}(0, \sigma^2), \tag{20}$$

while identifiability is enforced by constraining the additive effects to sum to zero and the multiplicative effects to have unit geometric mean across batches. These constraints ensure identifiability of the biological signal and batch parameters. Under this model, posterior inference is carried out jointly over all parameters,

$$P\left(\bar{F}, a_r, b_r, \sigma \mid \tilde{F}\right) \propto P\left(\tilde{F} \mid \bar{F}, a_r, b_r, \sigma\right) P(\bar{F}) P(a_r) P(b_r) P(\sigma), \tag{21}$$

yielding posterior distributions for the biological landscape and batch parameters. Batch-corrected measurements are obtained by inverting the measurement model using the posterior mean estimates of the batch parameters,

$$\tilde{F}_r^{\mathrm{adj}}(\mathbf{x}) = \hat{F}(\mathbf{x}) + \frac{\tilde{F}_r(\mathbf{x}) - \hat{F}(\mathbf{x}) - \hat{a}_r}{\hat{b}_r} \equiv \hat{F}(\mathbf{x}) + \xi_r(\mathbf{x}), \tag{22}$$

where $\xi_r(\mathbf{x})$ denotes the residual noise after removal of additive and multiplicative distortions, and we denoted posterior mean estimates by a hat, i.e., $\hat{F}(\mathbf{x}) = \mathbb{E}[\bar{F}(\mathbf{x}) \mid \tilde{F}]$, $\hat{a}_r = \mathbb{E}[a_r \mid \tilde{F}]$, and $\hat{b}_r = \mathbb{E}[b_r \mid \tilde{F}]$. Because $\xi_r(\mathbf{x})$ is a plug-in residual constructed from estimated parameters, it should not be identified with the generative noise $\pi_r(\mathbf{x})$ and generally departs from the assumed Gaussian *i.i.d.* model due to uncertainty propagation. Finally, the adjusted replicate maps $\tilde{F}_r^{\mathrm{adj}}(\mathbf{x})$ are therefore used only to characterize residual variability and quantify uncertainty, while all reported interaction measures are computed from the point estimate $\hat{F}(\mathbf{x})$.

The batch-removal procedure was validated on synthetic landscapes with known ground truth, where controlled additive and multiplicative batch effects were introduced. After correction, the inferred biological landscape accurately recovered the true signal without introducing spurious structure (see Figs. 2 and 3).

*Link functions for strictly positive landscapes*

Some biological traits, such as biomass or abundances, are strictly positive and exhibit multiplicative variability. In such cases, it is advantageous to apply a monotone link function $g$ and perform inference on the transformed measurements $Y_r(\mathbf{x}) = g(\tilde{F}_r(\mathbf{x}))$. For instance, a logarithmic link, $g(\cdot) = \log(\cdot)$, yields a Gaussian model in log-space –equivalent to a log-normal error model in the original scale–, while ensuring that the batch-corrected landscape remains strictly positive in the original scale.

In our implementation, to handle configurations with null function values when using a logarithmic link, we introduce a small positive offset $\epsilon \ll 1$ (typically $10^{-3}$), such that measurements are transformed as $Y_r(\mathbf{x}) = \log(\tilde{F}_r(\mathbf{x}) + \epsilon)$. This ensures numerical stability by mapping biological zeros to finite values in log-space. Additionally, for ecological consistency, the framework allows the all-zero community ($\mathbf{x} = \mathbf{0}$) to be excluded from the adjustment when it is consistently null across replicates, preventing the introduction of spurious biological signals in non-viable configurations.

#### S4 Propagation of measurement noise to interaction measures

All derived quantities considered in this work —including local epistatic interactions, Walsh–Hadamard coefficients, order-specific variances, and epistasis amplitudes— are computed from the replicate-aggregated landscape $\hat{F}(\mathbf{x})$. When multiple measurements are available for a given configuration, we define

$$\hat{F}(\mathbf{x}) = \frac{1}{R(\mathbf{x})} \sum_{r=1}^{R(\mathbf{x})} \tilde{F}_r^{\mathrm{adj}}(\mathbf{x}), \tag{23}$$

which provides an unbiased estimator of the underlying biological signal at that configuration. Unequal replication therefore affects only the uncertainty of the inferred quantities, not the geometric structure of the Walsh–Hadamard expansion, provided that the full configuration space is observed.

After batch correction, the residual term $\xi_r(\mathbf{x})$ is treated as a zero-mean random fluctuation, $\langle \xi_r(\mathbf{x}) \rangle_R = 0$, since any systematic bias can be absorbed into the definition of the biological landscape $\bar{F}(\mathbf{x})$. We further assume that residual noise is uncorrelated across configurations —an assumption that will be relaxed in SI S7—,

$$\mathrm{Cov}[\xi_r(\mathbf{x}), \xi_r(\mathbf{x}')] = 0 \qquad \text{for } \mathbf{x} \neq \mathbf{x}', \tag{24}$$

while allowing for heteroscedasticity, i.e., a configuration-dependent noise variance $\mathrm{Var}[\xi_r(\mathbf{x})]$. The effect of correlated residuals is discussed separately.

##### S4.1. Epistatic interactions

A local epistatic interaction of order $k$ is defined as an alternating sum over $2^k$ community configurations, as given in Eq. (3). When evaluated on noisy measurements, this quantity combines noise contributions evaluated at $2^k$ distinct configurations of the community. Because the residual noise has zero mean, the expectation value of the noise contribution vanishes. However, its variance does not cancel. Under the assumptions stated above, the noise contribution to a $k$-way interaction associated with the set of species $\mathbf{s}$ in a fixed background $\mathbf{B}$ has variance given by

$$\mathrm{Var}\left[\varepsilon_{\mathbf{s}}^{\mathrm{noise}}\right] = \sum_{U \subseteq S} \mathrm{Var}\left[\xi\left(\mathbf{B} + \mathbf{u}\right)\right], \tag{25}$$

that is, the sum of the noise variances associated with the $2^k$ configurations entering the interaction. In the homoscedastic case, $\mathrm{Var}[\xi(\mathbf{x})] = \sigma_\xi^2$, this expression simplifies to

$$\mathrm{Var}\left[\varepsilon_{\mathbf{s}}^{\mathrm{noise}}\right] = 2^k \sigma_\xi^2. \tag{26}$$

This exponential amplification reflects the combinatorial structure of higher-order interactions and arises even under the most favorable assumptions of independent measurement noise. Allowing for heteroscedasticity does not alter this conclusion and may further worsen the signal-to-noise ratio. As a result, the detectability of epistatic interactions deteriorates rapidly with interaction order.

##### S4.2. Noise projection on the Walsh–Hadamard coefficients

We now analyze how residual measurement noise propagates to the WH coefficients of the landscape. Because the WH transform is a linear change of basis, the noise contribution to each coefficient is obtained by projecting the residual term $\xi_r(\mathbf{x})$ onto the corresponding basis function. Writing the measured landscape as $\tilde{F}(\mathbf{x}) = \bar{F}(\mathbf{x}) + \xi_r(\mathbf{x})$, the WH coefficient associated with subset $\mathbf{s}$ can be decomposed as

$$f_{\mathbf{s}} = \bar{f}_{\mathbf{s}} + \eta_{\mathbf{s}}, \tag{27}$$

where $\bar{f}_{\mathbf{s}}$ denotes the coefficient of the underlying biological landscape and

$$\eta_{\mathbf{s}} = 2^{-N} \sum_{\mathbf{x}} \phi_{\mathbf{s}}(\mathbf{x}) \xi_r(\mathbf{x}) \tag{28}$$

is the noise contribution. For clarity, replica indices are omitted for $f_{\mathbf{s},r}/\eta_{\mathbf{s},r}$ in this section, as we focus on the typical noise propagation for a single measurement. Under the assumptions of zero-mean residuals and independence across configurations, the expectation of $\eta_{\mathbf{s}}$ vanishes. Its variance is given by

$$\text{Var}[\eta_{\mathbf{s}}] = 2^{-2N} \sum_{\mathbf{x}} \text{Var}[\xi(\mathbf{x})], \tag{29}$$

that is, by the average of the residual noise variances over all configurations. In the homoscedastic case $\text{Var}[\xi(\mathbf{x})] = \sigma_\xi^2$, this reduces to

$$\text{Var}[\eta_{\mathbf{s}}] = 2^{-N} \sigma_\xi^2. \tag{30}$$

Although the noise variance projected onto each individual WH coefficient is small and independent of the interaction order, the number of coefficients associated with a given order $k$ grows combinatorially as $\binom{N}{k}$. Consequently, as shown in the next section, when these coefficients are aggregated to estimate the total functional variance at order $k$, the cumulative noise contribution scales with the number of terms.

##### S4.3. Contribution of noise to order-specific variances

We now analyze how the noise projected onto the Walsh–Hadamard coefficients contributes to the variance associated with a given interaction order. For a fixed order $k$, the empirical variance is obtained by summing the squared coefficients over all subsets $\mathbf{s}$ of size $|\mathbf{s}| = k$,

$$\tilde{V}(k) = \sum_{\substack{\mathbf{s} \in \{0,1\}^N: \\ |\mathbf{s}|=k}} \left(\bar{f}_{\mathbf{s}} + \eta_{\mathbf{s}}\right)^2. \tag{31}$$

For $k > 0$, the projected noise has zero mean, $\langle \eta_{\mathbf{s}} \rangle_R = 0$, and the cross terms vanish in expectation. The expected value of the noisy variance estimate therefore reads

$$\langle \tilde{V}(k) \rangle_R = \bar{V}(k) + \sum_{\substack{\mathbf{s} \in \{0,1\}^N: \\ |\mathbf{s}|=k}} \text{Var}\left[\eta_{\mathbf{s}}\right], \tag{32}$$

where $\bar{V}(k) = \sum_{|\mathbf{s}|=k} \bar{f}_{\mathbf{s}}^2$ denotes the true biological variance at order $k$. Under the assumption of independent residual noise across configurations, all non-constant WH modes have comparable noise variance. Denoting this variance by $\text{Var}[\eta_{\mathbf{s}}]$, the noise contribution to the order-$k$ variance becomes

$$\left\langle \tilde{V}(k) \right\rangle_R = \bar{V}(k) + \binom{N}{k} \text{Var}[\eta_{\mathbf{s}}]. \tag{33}$$

#### S5 Uncertainty quantification and limits of detectability

Although the Bayesian batch-correction model (Section S2) provides the optimal point-estimate for the underlying biological landscape $\bar{F}(\mathbf{x})$, its intrinsic parametric uncertainty is unreliable for quantifying the final interaction measures. The use of a robust, non-parametric method is necessitated by two critical issues:

- **Residuals are Complex and Heteroscedastic:** The residual noise $\xi_r(\mathbf{x})$ (defined in S2) is a plug-in quantity derived from posterior estimates. This derivation causes the residuals to depart from the generative model's parametric assumptions, often resulting in high **heteroscedasticity** (variance dependent on the background configuration).
- **Metrics are Highly Non-Linear:** The statistics of ultimate interest, such as the order-specific variances $V(k)$ (Section S1.D) and higher-order epistasis, are highly non-linear functions of the landscape. Propagating analytical posterior uncertainty through these complex transformations is unreliable and computationally intractable.

For these reasons, uncertainty and detectability are robustly quantified using a resampling method, applied directly to the batch-corrected data. This non-parametric approach reliably captures the true sampling distribution of our final estimators.

##### S5.1. Bootstrap-based uncertainty estimation

Uncertainty in interaction measures is quantified using a non-parametric bootstrap applied to the batch-corrected landscapes. Because the distribution of the residual noise $\xi_r(\mathbf{x})$ is generally unknown and may exhibit heteroscedasticity or weak correlations, we avoid parametric modeling assumptions. Bootstrap samples are generated at the replicate level from the empirical residual maps $\xi_r(\mathbf{x})$ obtained after batch correction. Specifically, synthetic noise realizations are constructed as

$$\beta^{(b)}(\mathbf{x}) = \frac{1}{\sqrt{R(R-1)}} \sum_{r=1}^{R} w_r^{(b)}\, \xi_r(\mathbf{x}), \tag{34}$$

where $w_r^{(b)}$ are i.i.d. random multipliers with zero mean and unit variance (e.g., Normal or Rademacher variables). Each bootstrap landscape is then obtained as $F^{(b)}(\mathbf{x}) = \hat{F}(\mathbf{x}) + \beta^{(b)}(\mathbf{x})$. This *wild-cluster* (multipliers) bootstrap preserves the empirical variance structure of the residuals and, when present, their correlation across configurations (see S7 for discussion and proof). In the absence of cross-configuration correlations, the procedure matches the i.i.d. noise scaling expected under standard bootstrap/parametric nulls, while remaining non-parametric and typically more conservative. Repeating this procedure yields empirical distributions for interaction statistics, from which confidence intervals and detectability thresholds are computed.

##### S5.2. Bootstrap null model

The same bootstrap framework also defines an operational null model for detectability. Under this null hypothesis, the biological landscape contains no structured interactions, and all apparent signals arise solely from measurement noise.

Null bootstrap landscapes are constructed by resampling the residuals $\xi_r(\mathbf{x})$ without reintroducing the estimated biological signal,

$$F_{\text{null}}^{(b)}(\mathbf{x}) = \beta^{(b)}(\mathbf{x}), \tag{35}$$

where $\beta^{(b)}(\mathbf{x})$ is defined as above. Applying the same analysis pipeline to these null maps yields empirical null distributions for all interaction statistics considered.

As a consistency check, we compared this wild-cluster bootstrap null with a parametric bootstrap in a controlled synthetic setting where the true noise distribution is known. As shown in Fig. 5, for a purely Gaussian i.i.d. landscape the wild bootstrap accurately reproduces the order-dependent null epistasis amplitude and confidence intervals obtained from the parametric bootstrap. The wild procedure yields slightly broader intervals, reflecting its more conservative nature, but captures the same noise-induced scaling in the WH basis.

##### S5.3. Bootstrap *p*-values for a generic statistic $Q(\mathbf{x})$

Detectability is assessed by comparing the observed value of a landscape-derived quantity $Q$–individual epistatic coefficients, WH coefficients, epistasis amplitude or order-specific variances–, to its distribution under the bootstrap null model. Let $Q_{\text{obs}}$ denote the observed statistic –like local epistasic interactions Eq. (3), WH coefficients Eq. (7), etc.–, and $\{Q^{(b)}\}_{b=1}^{B}$ the corresponding values obtained from the null bootstrap maps $\{F_{\text{null}}^{(b)}\}$.

A two-sided non-parametric *p*-value is computed as

$$p = \frac{1}{B} \sum_{b=1}^{B} \mathbf{1}\Big\{ \big|Q^{(b)}\big| \geq \big|Q_{\text{obs}}\big| \Big\}, \tag{36}$$

which measures how often the noise-only null model produces a value at least as extreme as the observed one.

For statistics defined across multiple backgrounds, the same procedure is applied independently to each background. Because the null distributions are generated from the empirical residuals, the resulting *p*-values automatically reflect heteroscedasticity and any residual correlations present in the data. When only few replicates are available, the null distributions tend to be wide, yielding conservative $p$-values that faithfully represent the limited statistical power of the experiment. This procedure is visualized in Figs. 6 and 7, which show volcano plots of bootstrap $p$-values for local epistatic coefficients and WH modes, respectively, resolved by interaction order for the empirical dataset analyzed in this work.

##### S5.4. Detectability limits for higher-order interactions

The bootstrap-based $p$-values provide an operational definition of detectability. At a more conceptual level, the same comparison can be interpreted in terms of a signal-to-noise ratio that highlights the factors limiting the detection of higher-order interactions. All interaction measures considered in this work are estimated from replicate-averaged landscapes. For a linear map-derived quantity $Q$ estimated as $\hat{Q}$ from $R$ replicates, the estimator variance scales as

$$\mathrm{Var}[\hat{Q}] = \frac{1}{R}\mathrm{Var}[\chi], \tag{37}$$

where $\chi$ denotes the noise propagated through the corresponding linear combination of measurements. Detectability requires that the underlying biological signal exceeds the typical fluctuations induced by measurement noise, which can be expressed as

$$\mathrm{SNR}(Q) = \frac{|\bar{Q}|}{\sqrt{\mathrm{Var}[\chi]/R}} \gtrsim 1. \tag{38}$$

This condition clarifies how detectability depends on both the number of replicates and the structure of the interaction measure. For higher-order interactions, the propagated noise variance grows rapidly with interaction order due to their combinatorial definition. In experimental settings, where the number of replicates is typically limited, this growth leads to a sharp practical bound on the highest interaction order that can be reliably detected.

#### S6 Analysis of previously published datasets

In addition to the empirical dataset generated in this work, we analyzed several previously published microbial community datasets spanning different ecological contexts and experimental designs. These include the bacterial biodiversity–ecosystem functioning experiment of Langenheder et al. (2010) [3], from which we focus on the community–function landscape corresponding to growth on glucose and xylose as carbon sources; the microbial colonization landscapes of Gorostiaga et al. (2019) [4]; the replicated microbial community-function landscapes of Díaz-Colunga et al. (2024) [2]; and the duckweed-associated ecological landscape dataset of Ishizawa et al. (2025) [5]. All datasets provide complete combinatorial coverage of species assemblages, with the exception of Gorostiaga et al. (2019), for which 54 out of the 64 possible configurations are available.For this dataset, WH coefficients were computed using the available configurations only. Because the full hypercube is not sampled, the WH basis is not orthogonal under the empirical support, and the resulting variance decomposition does not correspond to an exact geometric partition. These results should therefore be interpreted with caution. Among all datasets, only Díaz-Colunga et al. (2024) includes replicate measurements.

Accordingly, uncertainty quantification and detectability were assessed using the bootstrap framework only for the Díaz-Colunga et al. dataset, while results for the remaining datasets are presented for qualitative comparison only. Figure 8 shows the distribution of absolute WH coefficients $|f_{\mathbf{s}}|$ as a function of interaction order. For Díaz-Colunga et al. (2024), replicate measurements allow estimation of a bootstrap-based null detection limit, enabling classification of coefficients as significant or nonsignificant. In contrast, for datasets lacking replication no uncertainty or detectability threshold can be inferred, and the corresponding coefficients should not be interpreted as statistically significant evidence of higher-order interactions. Across all datasets, including the one generated in this work, the typical magnitude of the WH coefficients decreases systematically with interaction order, indicating that the underlying biological signal is dominated by low-order interactions.

Despite these differences in experimental design, the order-resolved variance decomposition (Fig. 9) exhibits a consistent qualitative structure across datasets (including our in Fig. 3 of the main text). In all cases, the variance contribution $V(k)$ decays with interaction order, reflecting the strong structural constraint imposed by the combinatorial factor $\binom{N}{k}4^{-k}$, which peaks at order $k = 1$. At the same time, the epistasis amplitude $\langle \mathcal{E}^2 \rangle_k$ increases at higher orders, consistent with the accumulation of projected measurement noise over the rapidly growing number of WH modes at each order.

#### S7 Extension to correlated residuals

In practice, residual correlations can arise even when the underlying measurement noise is independent, as the residuals obtained after batch correction are plug-in quantities derived from estimated parameters. Shared uncertainty in the inferred biological landscape, identifiability constraints on batch effects, and the small number of experimental replicates can all induce weak cross-configuration correlations that are purely statistical in origin. These correlations should not be interpreted as genuine environmental or biological coupling between configurations.

This behavior is illustrated in Fig. 4. Before batch correction, measurements corresponding to different configurations are strongly correlated across replicates, reflecting shared batch-level fluctuations. After batch correction, this global correlation structure is removed and the distribution of pairwise correlations becomes symmetric and centered around zero, consistent with finite-sample noise.

In the empirical datasets analyzed here, replicate-level residual correlations across configurations are weak and largely compatible with finite-sample effects. Nevertheless, we include the correlated-residual bootstrap for methodological completeness and robustness; its inclusion does not qualitatively affect any of the results reported.

*a. Correlated noise in local epistasis.* When residuals are correlated, the propagation of noise to interaction measures is modified but not fundamentally altered. A local epistatic interaction of order $k$ is an alternating sum over $2^k$ configurations, Eq. (3). Writing the noise contribution as

$$\varepsilon_{\mathbf{s}}^{\text{noise}}(\mathbf{B}) = \sum_{U \subseteq S} (-1)^{k-|U|} \xi(\mathbf{B} + \mathbf{u}) \,, \tag{39}$$

where $U$ and $V$ denote subsets of the focal species set $S$, and $\mathbf{u}$ and $\mathbf{v}$ their corresponding indicator vectors, the variance of the noise contribution reads

$$\text{Var}\left[\varepsilon_{\mathbf{s}}^{\text{noise}}\right] = \sum_{U \subseteq S} \text{Var}[\xi(\mathbf{B} + \mathbf{u})] + 2 \sum_{\substack{U,V \subseteq S \\ U < V}} (-1)^{|U|+|V|} \text{Cov}[\xi(\mathbf{B} + \mathbf{u}) \,, \xi(\mathbf{B} + \mathbf{v})] \,. \tag{40}$$

Thus, correlations can change the effective noise prefactor and may introduce order-dependent effects, but they do not generically remove the combinatorial growth associated with summing $2^k$ terms. Positive correlations typically inflate the noise level of higher-order interactions, whereas systematic cancellations would require highly fine-tuned alternating correlation patterns.

*b. Noise projection onto Walsh–Hadamard modes.* A similar conclusion holds for WH coefficients. Writing the residual noise across configurations as a vector $\boldsymbol{\xi}$ and denoting by $\mathbf{C}$ its covariance matrix, the noise projected onto the WH basis has a correlator given by

$$\langle \eta_{\mathbf{s}} \eta_{\mathbf{s}'} \rangle_R = 2^{-2N} \, \phi_{\mathbf{s}}^\top \mathbf{C} \phi_{\mathbf{s}'} . \tag{41}$$

Correlations therefore induce non-uniform noise levels across WH modes, but do not eliminate the averaging over backgrounds that makes WH coefficients far more stable than individual local epistatic coefficients at the same order.

In practice, $\mathbf{C}$ is unknown and must be estimated from replicate-level residuals. Stacking residuals $\xi_r(\mathbf{x})$ as columns of the matrix $\boldsymbol{\Xi} = [\xi_1(\mathbf{x}), \ldots \xi_R(\mathbf{x})] \in \mathbb{R}^{2^N \times R}$ yields the sample covariance estimator

$$\hat{\mathbf{C}} = \frac{1}{R-1} \boldsymbol{\Xi} \boldsymbol{\Xi}^\top . \tag{42}$$

Because $R$ is often small, $\hat{\mathbf{C}}$ can be noisy, and some apparent correlation structure may reflect finite-sample effects rather than reproducible cross-configuration coupling.

*c. Propagation to order-resolved variances.* Correlated noise also affects the variance decomposition by interaction order. Let $\tilde{V}(k)$ be the empirical variance contribution at order $k$ computed from noisy WH coefficients. For $k \geq 1$, the expected noisy estimate decomposes into the biological signal plus a noise term,

$$\left\langle \tilde{V}(k) \right\rangle_R = \sum_{\substack{\mathbf{s}: \\ |\mathbf{s}|=k}} \bar{f}_{\mathbf{s}}^2 + \sum_{\substack{\mathbf{s}: \\ |\mathbf{s}|=k}} \text{Var}[\eta_{\mathbf{s}}]. \tag{43}$$

We summarize the average projected noise level at order $k$ by

$$K_k = \binom{N}{k}^{-1} \sum_{\substack{\mathbf{s}: \\ |\mathbf{s}|=k}} \text{Var}[\eta_{\mathbf{s}}] = \binom{N}{k}^{-1} 2^{-2N} \sum_{\substack{\mathbf{s}: \\ |\mathbf{s}|=k}} \phi_{\mathbf{s}}^\top \frac{\hat{\mathbf{C}}}{R} \phi_{\mathbf{s}}, \tag{44}$$

so that

$$\left\langle \tilde{V}(k) \right\rangle_R = \bar{V}(k) + \binom{N}{k} K_k . \tag{45}$$

Correlations therefore act by raising an order-dependent noise floor, which further limits the detectability of weak signals, especially at high orders.

*d. Wild-cluster (multiplier) bootstrap with correlated residuals.* Importantly, no additional modeling assumptions are required to incorporate correlated residuals within the bootstrap framework used in this work. The wild-cluster bootstrap, Eq. (34), constructs synthetic perturbations as linear combinations of replicate residuals, $\boldsymbol{\beta}^{(b)} \propto \boldsymbol{\Xi}\mathbf{w}$, and their covariance matches that of the empirical residuals

$$\langle \boldsymbol{\beta}^{(b)} \boldsymbol{\beta}^{(b)\top} \rangle_R = \frac{1}{R(R-1)} \boldsymbol{\Xi} \langle \mathbf{w}\mathbf{w}^\top \rangle_R \boldsymbol{\Xi}^\top = \frac{1}{R(R-1)} \boldsymbol{\Xi} I_R \boldsymbol{\Xi}^\top = \frac{1}{R} \hat{\mathbf{C}}. \tag{46}$$

Therefore, when correlations are present in the residuals, the bootstrap null inherits them automatically, yielding conservative uncertainty estimates and detectability assessments. A practical limitation is that with few replicates the estimate $\hat{\mathbf{C}}$ can be unstable, leading to wide null distributions and hence conservative $p$-values and confidence intervals; this correctly reflects the limited statistical power available in low-replicate landscapes.

Taken together, these considerations show that allowing for correlated residuals does not alleviate the detectability limits identified above. If anything, correlations typically inflate the effective noise level and further restrict the range of interaction orders that can be reliably resolved, reinforcing the conclusion that the scarcity of detectable high-order epistasis in empirical landscapes is not an artifact of overly simplified noise assumptions.

*e. Correlated residuals do not remove the exponential noise scaling.* When residual errors are correlated across configurations, the noise contribution to a $k$-way local epistatic interaction no longer comes only from summing $2^k$ variances: it also includes covariances between the $2^k$ terms entering the alternating sum, see Eq. (40). To suppress the exponential growth with $2^k$, the covariance term would need to cancel the variance term with comparable magnitude for all $k$. This would require a highly structured pattern of alternating positive and negative correlations that matches the sign pattern of the epistasis sum across all pairs of configurations.

In realistic settings, correlations are expected to vary smoothly with configuration similarity (e.g., decay with Hamming distance) and the residual covariance matrix must remain positive semidefinite. Under these mild and standard constraints, the covariance contributions cannot be tuned to track the rapidly oscillating combinatorial weights of the $k$-way epistasis operator. As a result, correlations may change the prefactor of the noise variance and make it mode- or order-dependent, but they do not eliminate the dominant combinatorial scaling with $2^k$. In practice, positive correlations typically increase the effective noise level of higher-order interactions, further reducing detectability; the bootstrap null model used in this work naturally incorporates these correlations and therefore provides a conservative data-driven estimate of uncertainty.

#### S8 Synthetic community landscapes

Synthetic functional maps provide a controlled benchmark to test and understand our statistical methods. Because the true generative model is known, they allow us to evaluate how well different approaches recover interactions, separate signal from noise, and identify potential failure modes. In this context, the epistasis amplitude $\{\langle \mathcal{E}_k^2 \rangle\}_{k=1,\dots,N}$, Eq. (12), offers a complete characterization of the global statistical structure of a synthetic map.

##### S8.1. Random maps

A random map represents the limiting case in which the biological signal is absent. In this scenario, measurements are generated purely from unstructured fluctuations across configurations,

$$F_r(\mathbf{x}) = a_r + b_r\, \epsilon_r(\mathbf{x}), \tag{47}$$

where $\epsilon_r(\mathbf{x})$ are i.i.d. random variables across $\mathbf{x}$ with zero mean and variance $\sigma_\epsilon^2$, and $a_r$ and $b_r$ capture potential additive and multiplicative batch effects. In our simulations we generate i.i.d. maps by drawing $U(\mathbf{x}) \sim \mathrm{Uniform}[a, b]$ independently for each $\mathbf{x}$ and then centering,

$$\epsilon(\mathbf{x}) = U(\mathbf{x}) - \frac{a+b}{2}, \tag{48}$$

so that $\langle \epsilon(\mathbf{x}) \rangle_{\mathbf{x}} = 0$ and $\sigma_\epsilon^2 = \frac{1}{12}(b-a)^2$. From Eq. (45), the expected variance spectrum reads

$$\big\langle \tilde{V}(k) \big\rangle_R = \binom{N}{k} 4^{-k} \langle \tilde{\mathcal{E}}_k^2 \rangle, \tag{49}$$

with $\langle \tilde{\mathcal{E}}_k^2 \rangle = 2^{2k-N} \sigma_\epsilon^2$. Other *i.i.d.* ensembles can be explored analogously: a Gaussian map $F(\mathbf{x}) \sim \mathcal{N}(0, \sigma^2)$ produces the same spectral form with $\sigma_\epsilon^2 = \sigma^2$, whereas a lognormal map yields a highly skewed distribution with a few extremely large values. This latter case is useful for studying the detection limits of high-order interactions, since only a small number of configurations dominate the amplitude at large $k$.

##### S8.2. Flat epistatic landscapes

To isolate the role of combinatorics from that of interaction strength, we also consider a class of synthetic landscapes with uniform epistatic amplitude across orders, which we refer to as *flat epistatic landscapes*. In these maps, all interaction orders contribute equally at the level of typical interaction strength, such that

$$\langle \mathcal{E}_k^2 \rangle = K^2 \qquad \text{for all } k \geq 1. \tag{50}$$

This condition is achieved by sampling Walsh–Hadamard coefficients with zero mean and order-dependent variance,

$$\mathrm{Var}(f_{\mathbf{s}}) = K^2\, 4^{-|\mathbf{s}|}, \tag{51}$$

so that the associated background-averaged epistatic interactions $\mathcal{E}_{\mathbf{s}} = 2^{|\mathbf{s}|} f_{\mathbf{s}}$ have constant variance across orders. In this construction, the total variance of the landscape reads

$$V_{\mathrm{tot}} = \sum_{k=1}^{N} V(k) = K^2 \left[ \left(\tfrac{5}{4}\right)^N - 1 \right], \tag{52}$$

which fixes the scale $K$ for a given target variance $V_{\mathrm{tot}}$. Flat epistatic landscapes therefore provide a useful reference case in which the epistasis amplitude does not decay with interaction order. Any structure observed in the variance spectrum $V(k)$ then arises purely from combinatorial and geometric factors, rather than from a hierarchy of interaction strengths. This makes them a natural benchmark to disentangle intrinsic epistatic structure from entropic effects.

In Fig. 10 we study the interplay between noise strength and replicability using a controlled synthetic experiment. Landscapes are constructed as the sum of a flat epistatic signal, $K^2 = 1$ taken as the ground-truth biological landscape, and independent Gaussian noise with standard deviation $\sigma \in \{0.5, 2, 5\}$. For each noise level we analyze replicate-averaged landscapes with $R \in \{3, 10, 20\}$. At fixed $\sigma$, increasing the number of replicates progressively suppresses noise-induced higher-order components, allowing the flat underlying signal to be recovered over an increasing range of interaction orders. Conversely, at fixed $R$, increasing noise strength inflates the apparent epistasis at high orders, closely following the noise-only null expectation. In the limit of small $\sigma$ and large $R$, the inferred epistasis amplitude converges to the true biological signal, indicating that deviations at finite $R$ arise from statistical detectability limits rather than genuine higher-order structure.

##### S8.3. Mechanistic testbed: generalized Lotka–Volterra landscapes

To validate our theory, we use the generalized Lotka–Volterra (gLV) model as a mechanistic testbed,

$$\frac{dy_i}{dt} = y_i \left( r_i + \sum_{j=1}^{N} a_{ij} y_j \right), \tag{53}$$

where $y_i$ denotes the abundance of species $i$, $r_i$ its intrinsic growth rate, and $a_{ij}$ the interaction matrix, with $a_{ii} = -1/K_i$ setting the carrying capacity. For a given community configuration $\mathbf{x}$, steady-state abundances $\{\bar{y}_i\}$ are obtained by solving the corresponding linear system. We define the community function as a linear combination of steady-state abundances,

$$F(\mathbf{x}) = \sum_{i=1}^{N} \varphi_i \bar{y}_i x_i, \tag{54}$$

where $\varphi_i$ is the biomass contribution of species $i$.

The interaction matrix $A = (a_{ij})$ is held fixed and sampled as follows. Self-interactions are fixed to $a_{ii} = -1$ for all species, setting the carrying capacity scale. Off-diagonal interactions $a_{ij}$ with $i \neq j$ are drawn independently from a Gaussian distribution with zero mean and variance $\sigma_a^2$. Only interaction matrices that yield feasible and stable coexistence equilibria are retained, i.e., those for which the steady state $\bar{\mathbf{y}} = -A^{-1}\mathbf{r}$ has strictly positive components [6] and is dynamically stable [7].

Crucially, although gLV dynamics only involve pairwise interactions in abundances, the steady-state landscape $F(\mathbf{x})$ can exhibit higher-order epistasis because the vector of equilibria $\bar{\mathbf{y}}$ depends on the inverse of the community sub-matrix. However, in our parameter regime, the noiseless gLV variance spectrum is effectively dominated by low-order terms, with contributions beyond second order being numerically negligible (Fig. 11A and E).

This allows us to isolate the effect of measurement noise. We now generate $R = 5$ synthetic experimental replicates by adding independent Gaussian noise $\pi_r(\mathbf{x}) \sim \mathcal{N}(0, \sigma^2)$ to the fixed underlying landscape. This induces an approximately order-independent noise variance per Walsh–Hadamard mode. When aggregated across the $\binom{N}{k}$ modes at order $k$, the noise in the replicate-averaged coefficients $\bar{\eta}_{\mathbf{s}}$ produces an apparent inflation of the higher-order variance $\tilde{V}(k)$ that follows the combinatorial scaling $\binom{N}{k}\mathrm{Var}[\bar{\eta}_{\mathbf{s}}]$ predicted by our theory (Eq. 33), even when the underlying mechanistic landscape contains no true signal at those orders, see Fig. 11.

### S9 Supplementary Tables

TABLE S1. Species-level taxonomic assignment of isolates

| Isolate(s) | Species |
|---|---|
| C1, C12 | *Dermacoccus nishinomiyaensis* |
| C4 | *Janthinobacterium agaricidamnosum* |
| C5 | *Micrococcus luteus* |
| C7 | *Arthrobacter* sp. Tibet-IX22 |
| C8, C9 | *Rhodococcus corynebacterioides* |
| C10 | *Arthrobacter agilis* |
| C14 | *Staphylococcus warneri* |

TABLE S2. Number of different SNPs

| Specie | C1 | C4 | C5 | C6 | C7 | C8 |
|---|---|---|---|---|---|---|
| C1 | - | 1062 | 1092 | 1042 | 334 | 1041 |
| C4 | 1062 | - | 545 | 414 | 1069 | 440 |
| C5 | 1092 | 545 | - | 262 | 1098 | 333 |
| C6 | 1042 | 414 | 262 | - | 1040 | 214 |
| C7 | 334 | 1069 | 1098 | 1040 | - | 1054 |
| C8 | 1041 | 440 | 333 | 214 | 1054 | - |
| C9 | 643 | 414 | 363 | 318 | 654 | 221 |
| C10 | 1067 | 437 | 283 | 167 | 1075 | 208 |
| C12 | 1057 | 421 | 286 | 144 | 1063 | 210 |
| C14 | 1054 | 376 | 505 | 372 | 1060 | 429 |

### S10 Supplementary Figures

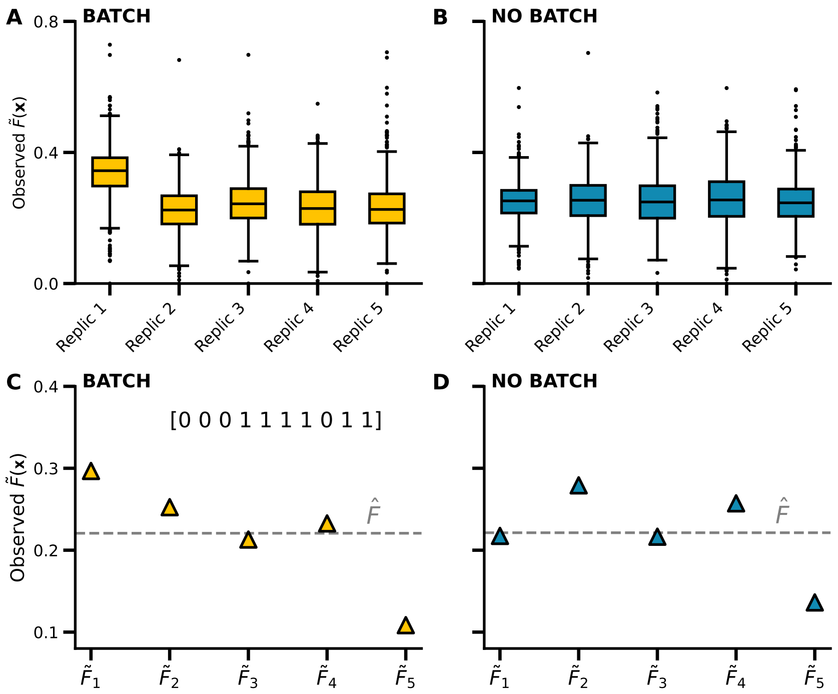


FIG. 1. **Example of batch effects and their correction**. **A**,**B.** Boxplots of the five experimental replicates of the observed function $\tilde{F}(\mathbf{x})$ (see SI. S2), with batch effects and after batch removal following the procedure described in SI S3 S3.2. Once the batch effect is removed, the boxes become aligned around a common central value. **C**,**D.** Observed values $\tilde{F}(\mathbf{x})$ for a specific species configuration, $\mathbf{x} = [0, 0, 0, 1, 1, 1, 1, 0, 1, 1]$, with batch effects C and without batch effects D. The dashed line indicates the estimated mean $\hat{F}$ shared across replicates after correction. Once the batch effect is removed, the distributions of the replicates align around a common central value. All subsequent analyses in this work use the batch-corrected data.

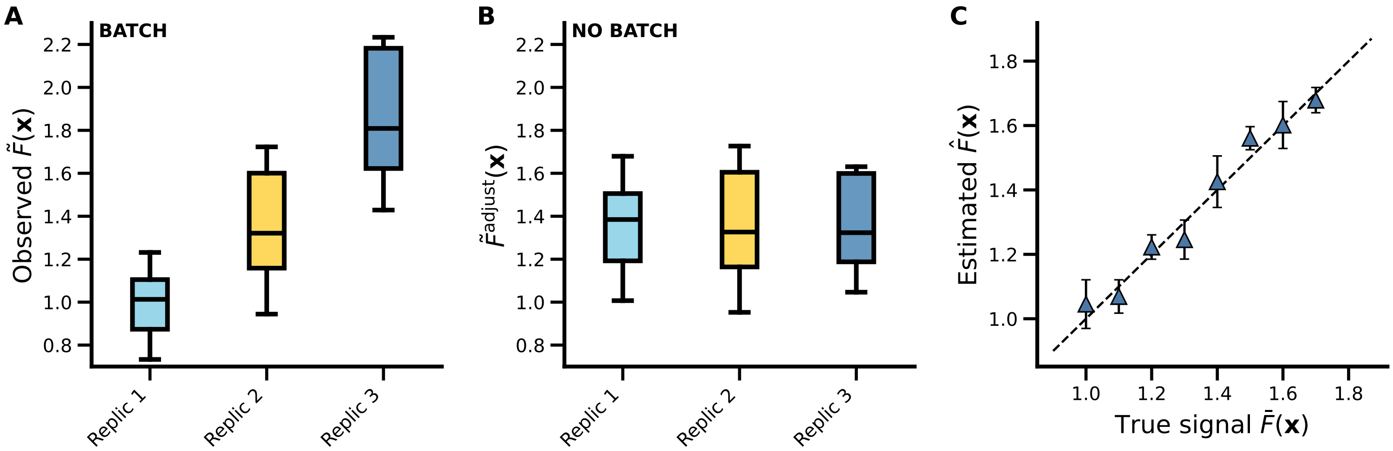


FIG. 2. **Batch effects, correction, and signal recovery in a synthetic dataset. A.** Observed measurements $\tilde{F}(\mathbf{x})$ for three synthetic replicates generated with known additive and multiplicative batch effects. Each box summarizes the distribution of $\tilde{F}(\mathbf{x})$ values across all configurations for a given replicate, revealing systematic shifts and scale differences induced by batch effects. **B.** The same replicates after batch-effect removal, $\tilde{F}^{\mathrm{adjust}}$. The distributions collapse onto a common scale, indicating successful correction and recovery of a shared underlying signal. **C.** Recovery of the underlying biological map. Each point represents a configuration $\mathbf{x}$, showing the batch-corrected estimate $\hat{F}(\mathbf{x})$ against the true value $\bar{F}(\mathbf{x})$. Vertical error bars indicate the standard deviation across replicates after correction. The corrected means lie tightly along the identity line (dashed), demonstrating accurate reconstruction of the true signal while preserving intrinsic biological variability.

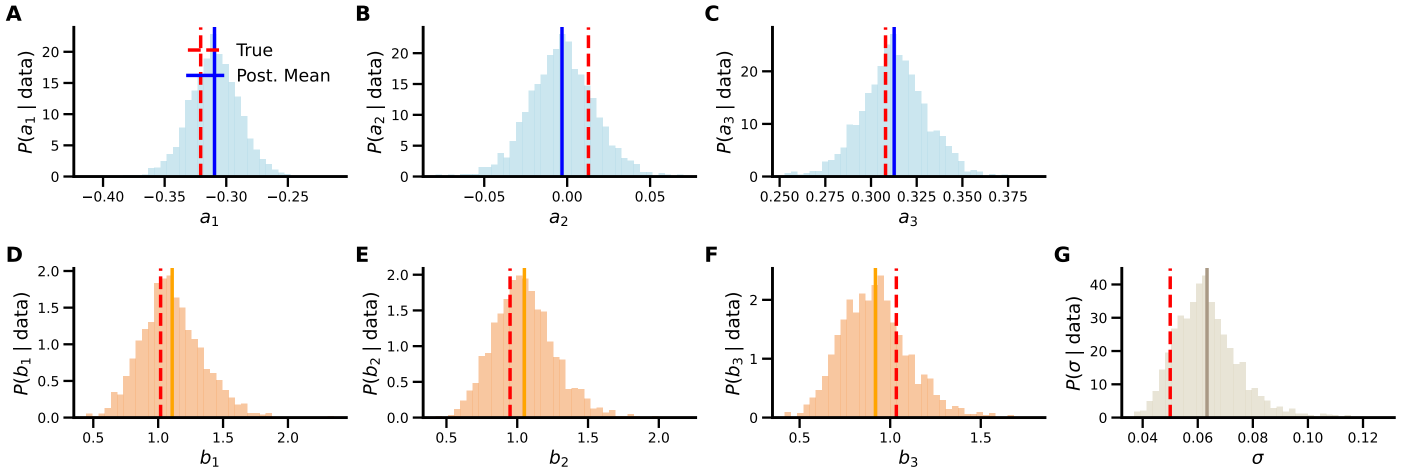


FIG. 3. **Posterior distributions of batch-effect parameters in the synthetic test.** Panels **A–C** show the marginal posterior distributions of the additive batch effects $a_r$, one panel per replicate, while panels **D–F** show the corresponding posteriors for the multiplicative batch effects $b_r$. Panel **G** displays the posterior distribution of the noise scale $\sigma$. In all cases, the posteriors are sharply peaked and span a narrow range, indicating low uncertainty in the inferred parameters. The posterior means (solid lines) lie very close to the true values used in the synthetic generative model (dashed red lines), demonstrating accurate and robust recovery of both additive and multiplicative batch effects, as well as the residual noise level.

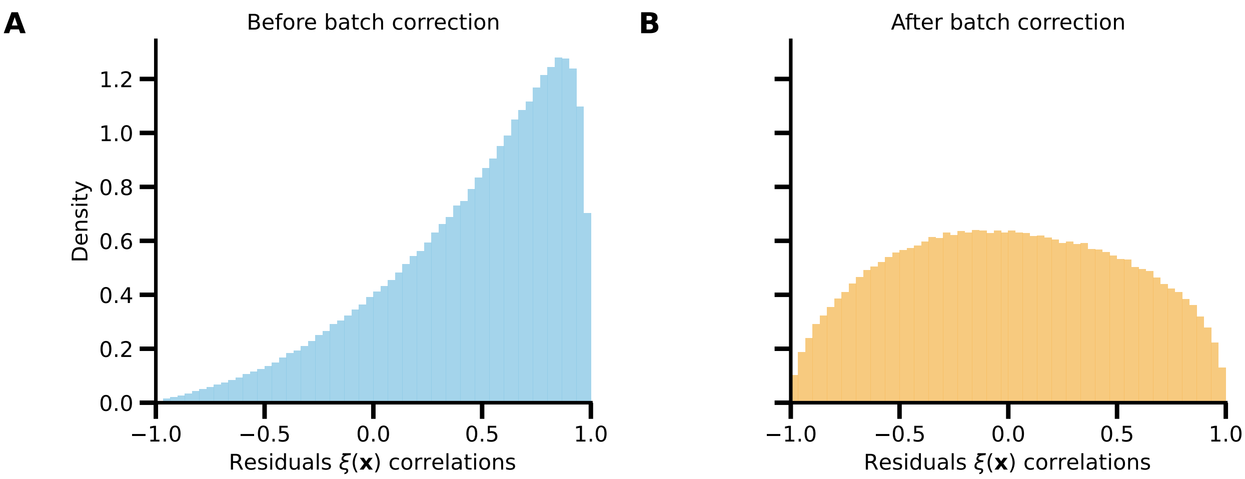


FIG. 4. **Residual correlations across community configurations before and after batch correction.** **A.** Distribution of pairwise Pearson correlations between community configurations computed across experimental replicates *before* batch correction. Strong positive correlations reflect shared batch-level fluctuations affecting all configurations within a replicate. **B.** The same distribution *after* batch correction. Global correlations are removed and the distribution becomes symmetric and centered around zero, consistent with finite-sample fluctuations of correlation estimates computed from a small number of replicates. This illustrates that residual cross-configuration correlations in the corrected data are weak and compatible with statistical noise rather than genuine biological coupling.

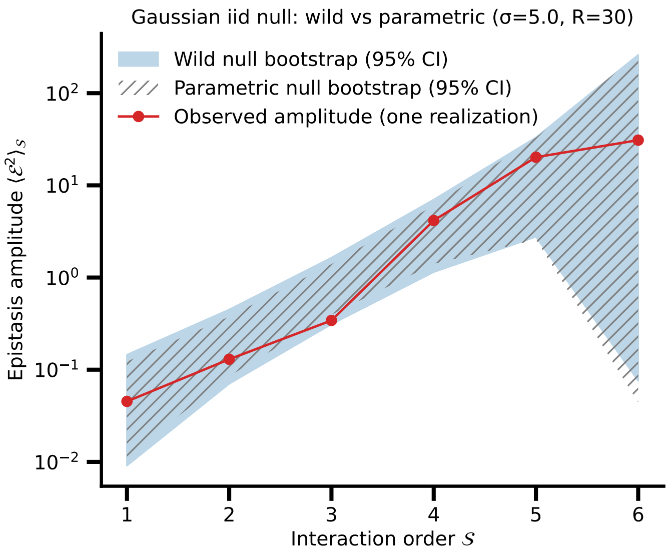


FIG. 5. Comparison between wild bootstrap and parametric bootstrap for a purely Gaussian i.i.d. synthetic landscape. Although the wild (multiplier-based) bootstrap does not assume the true underlying noise distribution, it recovers the same order-dependent null epistasis amplitude and produces confidence intervals consistent with those obtained from the parametric bootstrap, which explicitly assumes a Gaussian model. The wild bootstrap yields slightly broader intervals, reflecting its more conservative nature. This test demonstrates that the wild procedure robustly captures noise propagation in the Walsh–Hadamard basis even when the noise distribution is misspecified. Parameters: number of species $N = 6$, number of replicates $R = 30$, bootstrap samples $B = 1000$, confidence level $95\%$, and noise $\xi(\mathbf{x}) \sim \mathcal{N}(0, 5)$.

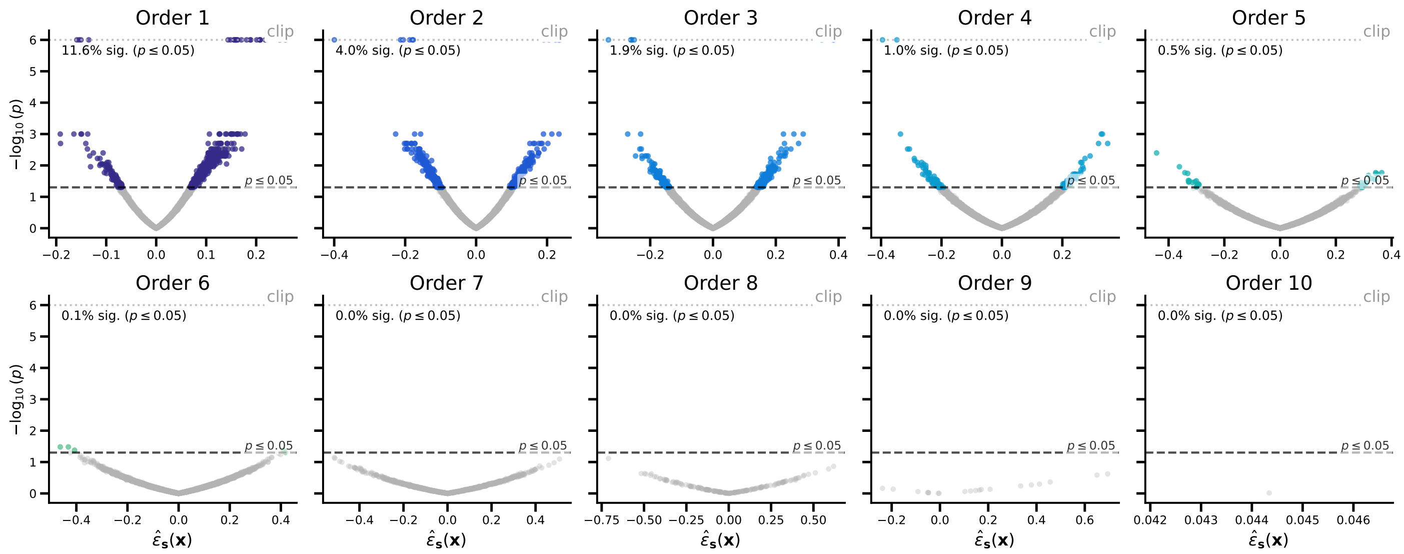


FIG. 6. **Volcano plots of local epistasis across interaction orders for empirical dataset.** Each panel shows a volcano plot for all coefficients of a given interaction order $k$, with epistasis $\hat{\varepsilon}_{\mathbf{s}}(\mathbf{x})$ on the $x$-axis and significance $-\log_{10}(p)$ on the $y$-axis. As interaction order increases, the strength of the epistasis coefficients tends to increase, while detection power diminishes due to broader null distributions. Beyond order 6, no coefficients are detected as significant. Notably, even at order 1 only about $10\%$ of coefficients surpass the detection threshold, consistent with the expectation that ecological interaction networks are sparse –most microbial pairs interacting only weakly and a minority exhibiting strong or functionally relevant interactions. $p$-values below numerical precision are clipped at $10^{-6}$ (dotted line), and the dashed horizontal line indicates the detection threshold $p \leq 0.05$. Percentages report the fraction of significant coefficients per order.

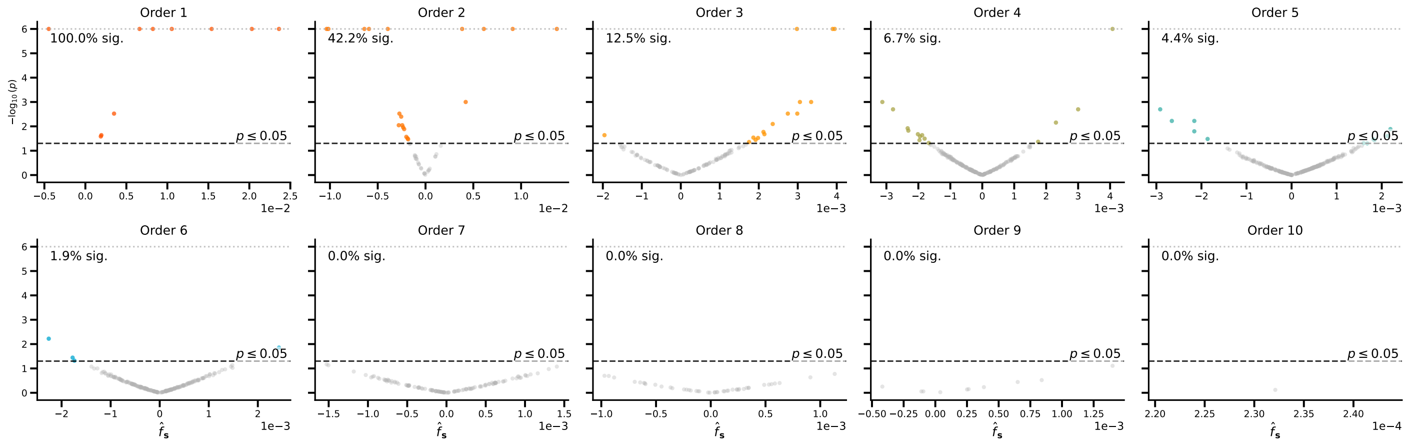


FIG. 7. **Volcano plots of Walsh–Hadamard coefficients across interaction orders for an empirical dataset.** Each panel shows a volcano plot for all Walsh–Hadamard modes of a given interaction order $k$, with the inferred coefficient $\hat{f}_{\mathbf{s}}$ on the $x$-axis and statistical significance $-\log_{10}(p)$ on the $y$-axis. As interaction order increases, the typical magnitude of the coefficients tends to decrease (see x-axis), while statistical power decreases as well due to it. Beyond order 6, no coefficients are detected as significant. $p$-values below numerical precision are clipped at $10^{-6}$ (dotted line), and the dashed horizontal line indicates the detection threshold $p \leq 0.05$. Percentages report the fraction of significant modes per interaction order.

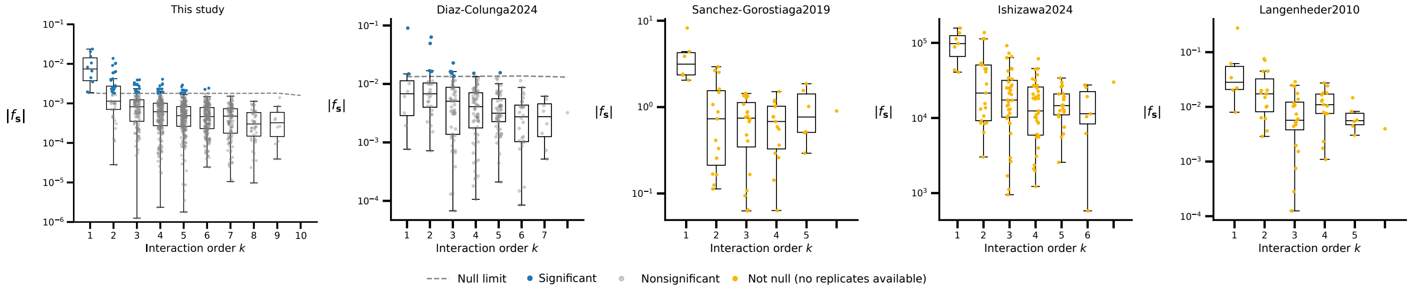


FIG. 8. Distribution of the absolute Walsh–Hadamard coefficients $|f_{\mathbf{s}}|$ as a function of interaction order $k$ for four experimental datasets. Each panel corresponds to a different study. For the empirical dataset generated in this work and for *Díaz-Colunga et al. (2024)*, replicate measurements allow estimation of uncertainty and a bootstrap-based null detection limit. Coefficients above the null limit (dashed line) are highlighted as significant, while coefficients below are classified as nonsignificant. For the remaining datasets, only a single measurement per configuration is available; therefore, neither uncertainty nor a null detection limit can be estimated. In these cases, coefficients are shown for qualitative comparison only and should not be interpreted as statistically significant.

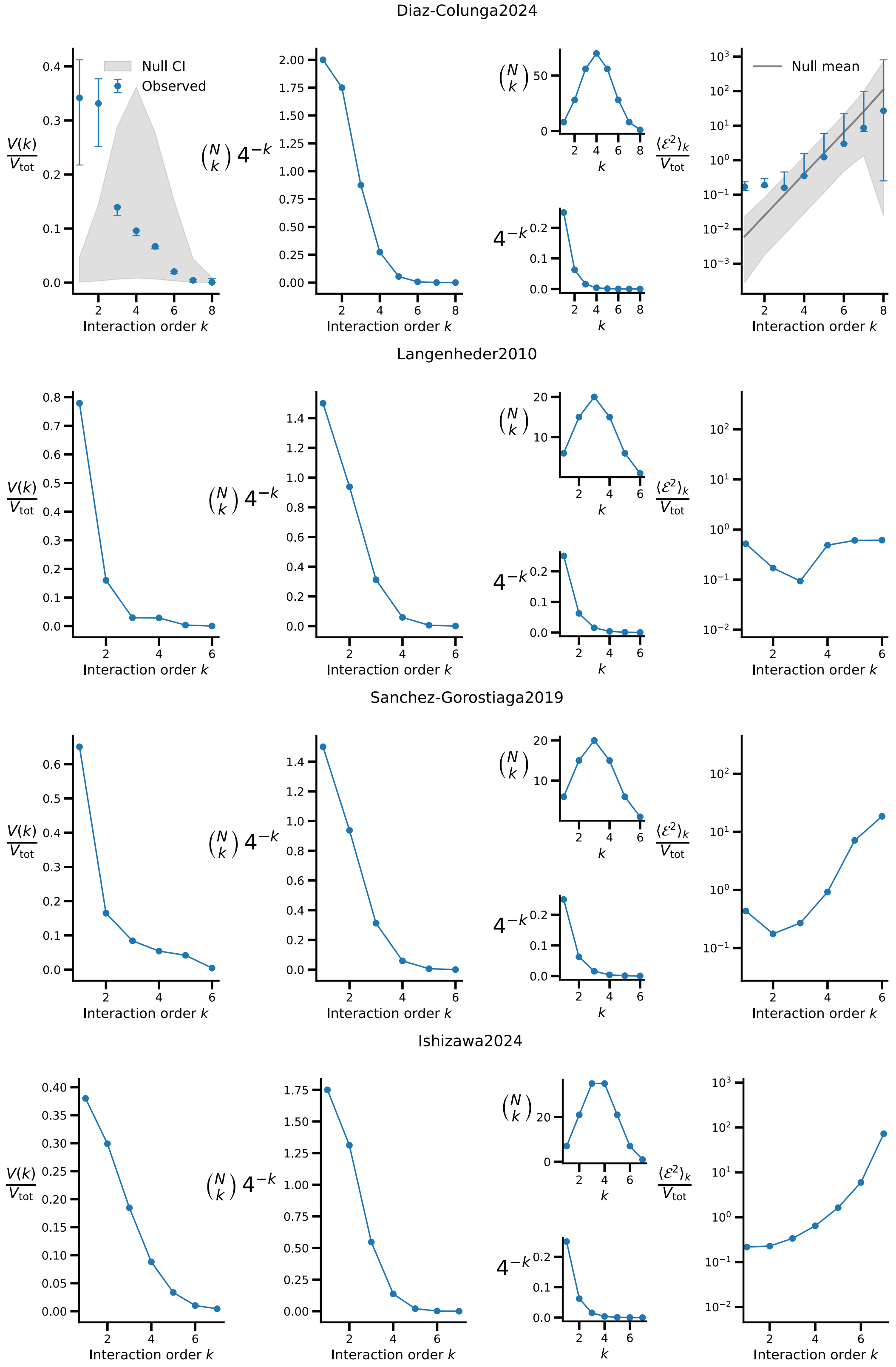


FIG. 9. **Order-resolved variance decomposition across experimental datasets.** Variance decomposition $V(k)/V_{\mathrm{tot}}$ (left panels), combinatorial weighting $\binom{N}{k}4^{-k}$ (second column), and epistasis amplitude $\langle\mathcal{E}^2\rangle_k/V_{\mathrm{tot}}$ (right panels) shown as a function of interaction order $k$ for four experimental datasets (one dataset per row). In all cases, the structural combinatorial factor $\binom{N}{k}4^{-k}$ peaks at $k = 1$, imposing a strong structural constraint on the contribution of higher-order interactions to the total variance. Across datasets, the variance spectrum $V(k)$ decays with interaction order, while the epistasis amplitude $\langle\mathcal{E}^2\rangle_k$ increases at high orders, consistent with the accumulation of projected measurement noise over the $\binom{N}{k}$ coefficients at each order. For Gorostiaga et al. (2019), the landscape is incomplete (54/64 configurations). The WH-based variance decomposition can not be interpreted as the contribution of each interaction order to the total variance, it is shown for qualitative comparison only. Moreover, only the dataset of Díaz-Colunga et al. (2024) includes replicate measurements, allowing estimation of uncertainty and a bootstrap-based null model; the remaining datasets lack replication and are therefore shown without null confidence intervals, for qualitative comparison.

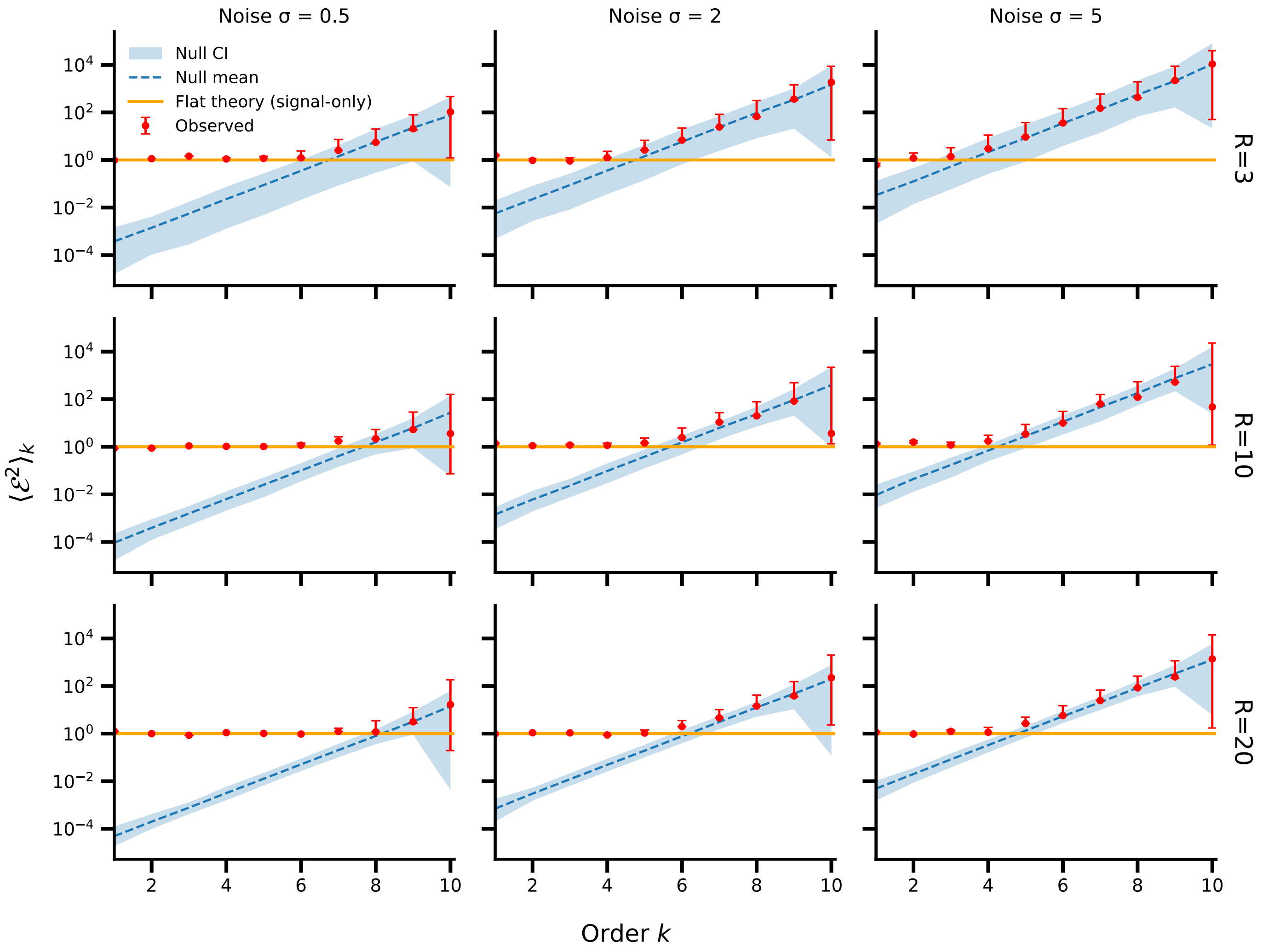


FIG. 10. **Recovery of a flat epistatic signal in the presence of measurement noise.** Epistasis amplitude $\langle \mathcal{E}^2 \rangle_k$ inferred from synthetic landscapes constructed as the sum of a flat epistatic signal with $K^2 = 1$ and independent Gaussian noise. Columns correspond to increasing noise strength $\sigma = 0.5, 2, 5$, while rows correspond to increasing numbers of replicates $R = 3, 10, 20$. Red points show the observed epistasis amplitude with bootstrap uncertainty bars. Blue shaded regions indicate the 95% confidence interval of the bootstrap null model, with the dashed line showing the null mean. The horizontal orange line denotes the flat epistatic amplitude of the underlying signal. At fixed noise level, increasing $R$ progressively suppresses noise-induced higher-order components, whereas at fixed $R$ increasing noise strength inflates apparent high-order epistasis in agreement with the null expectation.

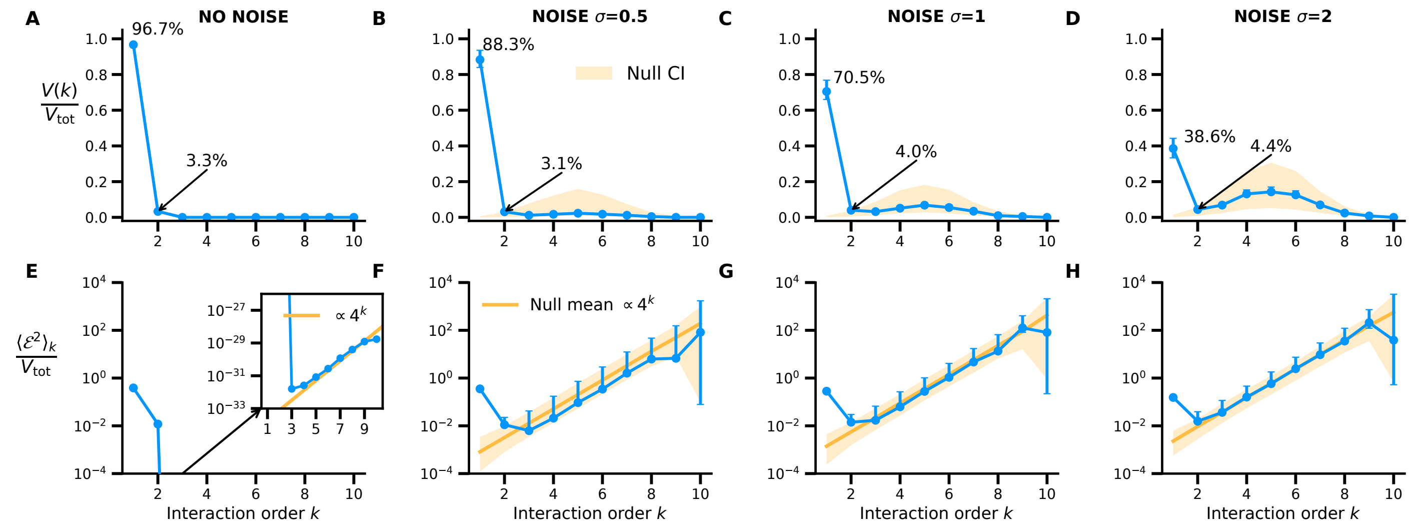


FIG. 11. **Emergence of noise-induced spurious higher-order interactions in mechanistic landscapes.** Community-function landscapes were simulated using a generalized Lotka–Volterra (gLV) model with $N = 15$ species. By construction, gLV steady states depend linearly on the abundances of present species (when feasible and stable). **(A–D)** Functional variance decomposition $V(k)$ (top row) and epistasis amplitude (bottom row) under increasing levels of Gaussian measurement noise $\sigma \in \{0, 0.5, 1, 2\}$. In the noiseless case **(A)**, the variance spectrum truncates abruptly after the second order, with values for $k \geq 3$ falling to the machine precision limit ($\sim 10^{-16}$, dotted line). As noise intensity increases **(B–D)**, the variance in higher orders is artificially inflated, following the combinatorial distribution $\binom{N}{k}$ predicted by our framework (Eq. 33). Note that while the epistasis amplitude (the typical strength of individual interactions, bottom row) remains at the noise-floor level for high orders, the total order variance becomes dominant at $k \approx N/2$ due to the accumulation of projected noise across the vast number of Walsh–Hadamard coefficients at those orders. These results confirm that apparent complexity in noisy landscapes can be a statistical artifact and validate our correction framework and detectability limits. Simulation parameters: connectance $c = 0.5$, off-diagonal interactions $a_{ij} \sim \mathcal{N}(0.2, 0.2)$, growth rates $r_i = 0.1$, and $R = 5$ synthetic replicates. Error bars indicate bootstrap confidence intervals across replicates, and shaded regions show the corresponding noise-only bootstrap null distributions; the yellow curve denotes the mean null epistasis amplitude scaling as $\propto 4^k$.
